# Analysis of motor-based transport in primary cilia by dynamic mode decomposition of live-cell imaging data

**DOI:** 10.64898/2026.03.27.714708

**Authors:** Fabiola Campestre, Line Lauritsen, Lotte B. Pedersen, Daniel Wüstner

**Affiliations:** Department of Biology, University of Copenhagen, Universitetsparken 13, DK-2100 Copenhagen Ø, Denmark; Department of Biochemistry and Molecular Biology, University of Southern Denmark, Campusvej 55, DK-5230 Odense M, Denmark

**Keywords:** KIF13B, intraflagellar transport, IFT172, super-resolution microscopy, dynamic mode decomposition, live-cell imaging

## Abstract

Kinesin-3 motor proteins are increasingly recognized for their important roles in cilia. The mammalian kinesin-3 motor KIF13B moves bidirectionally in primary cilia and regulates ciliary content, but its relationship to the intraflagellar transport (IFT) machinery is unclear. Here, we combine quantitative live-cell imaging with a new kymograph analysis based on dynamic mode decomposition (DMD) to separate mobile from immobile protein populations in primary cilia. This approach simplifies extraction of molecular velocities from kymographs and reveals that a KIF13B deletion mutant retaining only the motor domain and part of the forkhead-associated domain does not alter steady-state IFT velocity or frequency. However, when retrograde dynein-2 function is inhibited by Ciliobrevin D, both anterograde and retrograde IFT velocities decrease in parental cells, as expected, but remain unchanged in KIF13B mutant cells. Structured illumination, confocal, and STED microscopy further show that KIF13B localizes to the ciliary membrane and concentrates at the periciliary membrane region and the centriolar subdistal appendages, below the distal appendage marker FBF1. Our improved kymograph approach provides new insight into KIF13B ciliary function and simplifies the quantitative analysis of ciliary protein transport.

## INTRODUCTION

Bidirectional intraflagellar transport (IFT) is required for proper assembly, maintenance, and function of almost all types of cilia, including primary cilia that function as cellular signalling hubs during vertebrate development and adult homeostasis (Anvarian et al., 2019). Kinesin-2 motors, initially purified from sea urchin egg cytosol, are the primary drivers of anterograde movement of IFT trains towards the ciliary tip, moving at a velocity of ∼0.4 µm/s in mammalian cells (Cole et al., 1993; Morthorst et al., 2018). Kinesin-2 motors comprise the heterotrimeric kinesin-2 complex (KIF3A/KIF3B/KAP in vertebrates), broadly required for ciliogenesis, and homodimeric kinesin-2 motors, such as OSM-3 in *Caenorhabditis elegans* and KIF17 in vertebrates, often serving specialized roles (Pedersen & Rosenbaum, 2008; Prevo et al., 2017). In *C. elegans* neuronal sensory cilia, heterotrimeric kinesin-2 (composed of KLP-11, KLP-20, and KAP-1) and homodimeric OSM-3 motors cooperate during anterograde IFT along the proximal segment of the axoneme, comprising doublet microtubules, before OSM-3 alone takes over for transport along the distal singlet region (Evans et al., 2006; Prevo et al., 2015; Snow et al., 2004). The two motors exhibit distinct velocities and cargo preferences, suggesting a division of labor that ensures efficient IFT and proper cilium assembly (Pedersen & Rosenbaum, 2008; Prevo et al., 2017). In vertebrates, homodimeric kinesin-2 motor KIF17 similarly localizes to the distal segment of photoreceptor and olfactory cilia, where it respectively regulates outer segment elongation and the selective transport of ciliary membrane proteins, including cyclic nucleotide–gated channels (Dishinger et al., 2010; Insinna et al., 2008; Jenkins et al., 2006).

Beyond kinesin-2 motors, other kinesin family members are found to operate within primary cilia, contributing to ciliary biogenesis, homeostasis, or function (Morthorst et al., 2018; Reilly & Benmerah, 2019). For example, the kinesin-3 motor KIF14 moves processively within primary cilia, and its depletion leads to upregulation of AURKA phosphorylation, impaired Smoothened translocation into cilia, and reduced frequency of IFT (Mikulenková et al., 2025; Pejskova et al., 2020). Other kinesin-3 members, such as *C. elegans* KLP-6, regulate neuronal sensory ciliary composition and extracellular vesicle release to facilitate inter-organismal communication during mating (Peden & Barr, 2005; Wang et al., 2014), a process also influenced by kinesin-2 motors (Clupper et al., 2022). KLP-6 moves within cilia independently of kinesin-2 motors, acting as a positive regulator of ciliary growth, and *klp-6* mutant worms display increased anterograde velocity, from ∼0.73 to ∼0.89 µm/s, of a subpopulation of OSM-3 motors, within cephalic male cilia (Morsci & Barr, 2011). We previously demonstrated that the mammalian kinesin-3 motor, KIF13B, localizes to primary cilia of immortalized retinal pigment epithelial (hTERT-RPE1) cells, where it displays burst-like bidirectional movement at speeds similar to those of conventional IFT (Juhl et al., 2023). Moreover, we showed that KIF13B regulates ciliary membrane protein content and signalling at least in part by promoting endocytic retrieval at the ciliary base while suppressing release of large extracellular vesicles from cilia (Rezi et al., 2025; Schou et al., 2017). Despite these advances, it remains unknown whether and how KIF13B modulates the canonical IFT machinery in primary cilia.

Quantitative analysis of IFT is based on the measurement of molecular velocities from kymographs, i.e. space-time plots extracted from live-cell video sequences. Many computational tools have been developed for kymograph analysis, but there is a lack of methods which dissect the complex multiscale dynamics observed in cilia into different modes. Dynamic mode decomposition (DMD) is a data-driven numerical method, which has been developed for analysis of complex spatiotemporal datasets originally in the field of fluid dynamics (Kutz et al., 2016; Schmid, 2010). DMD is based on the singular value decomposition (SVD) of a space-time matrix, which is exactly the way, dynamic data is arranged in kymographs. DMD aims for a decomposition of dynamic data into spatial and temporal modes of differing frequencies which describe the progression of the time series. Slow modes can be identified with background in video sequences, for example, while the fast modes resemble image foreground (Erichson et al., 2019; Grosek & Kutz, 2014). DMD is finding many applications in biomedical imaging, including magnetic resonance imaging and positron emission tomography data, particularly in neuroscience (Casorso et al., 2019; Fu et al., 2020; Kunert-Graf et al., 2019; Tirunagari et al., 2017, 2019). We recently showed the potential of DMD for bleaching-based image segmentation, image denoising and analysis of 3D and polarimetry data in fluorescence microscopy (Wustner et al., 2024; Wüstner, 2022b). We also found DMD able to dissect dynamic live-cell time-lapse microscopy data, where either exponentially decaying intensities, such as in fluorescence loss in photobleaching experiments, or oscillating fluorescence signals, as observed in glucose-induced oscillations of autofluorescence in yeast, could be dissected in individual components, enabling in-depth analysis of the underlying dynamics (Wustner et al., 2025; Wüstner, 2022a).

Here, we demonstrate that DMD combined with time-delay embedding (DMD-TDE) can decompose the complex protein dynamics inside cilia into stationary and mobile populations, which dramatically enhances the accuracy and speed of automated kymograph analysis routines. We show that DMD-TDE but not standard DMD allows for accurate reconstruction of synthetic and experimental kymographs, and we demonstrate that this is possible because DMD-TDE approximates the underlying Koopman operator with time-delays of the data as observables. Using live-cell imaging combined with DMD-TDE-based kymograph analysis, we find that a deletion mutant in KIF13B, retaining only the motor domain and part of the forkhead-associated (FHA) domain, does not significantly affect the conventional anterograde and retrograde IFT velocities within primary cilia of untreated hTERT-RPE1 cells. When cells are treated with the dynein inhibitor Ciliobrevin D, which inhibits retrograde IFT in wild-type cells, retrograde IFT in KIF13B mutant cells appears unaffected. Halo-KIF13B kymograph tracks show a rapid exit from the cilium that fits well with a combination of diffusion and active retrograde transport. Thanks to super-resolution imaging, we are able to visualize intraciliary KIF13B localizing in proximity to IFT172, but closer to the ciliary membrane than to the axoneme. Additionally, KIF13B was concentrated at the periciliary membrane region and the centriolar subdistal appendages, below the distal appendage marker FBF1. These observations support a model whereby KIF13B functions as a scaffold for protein recruitment to promote endocytic retrieval and microtubule plus end-directed endolysosomal trafficking of ciliary components away from the ciliary base. Overall, our findings indicate that the loss of KIF13B has little impact on conventional IFT transport rates in primary cilia and that KIF13B primarily influences ciliary protein content through an IFT-independent mechanism.

## RESULTS AND DISCUSSION

### Dynamic mode decomposition of synthetic and experimental kymograph data

Kymographs can be considered as discrete snapshots of the system in time, and the aim of DMD is to approximate the evolution operator, *A*, which describes the progression of the system from one snapshot to the next (see Materials and Methods). For that, DMD uses the SVD of the data matrix which allows for spectral decomposition of *A* and thereby enables one to identify the dominant modes describing the observed dynamics. A low-rank approximation captures only slowly evolving processes, while large displacements of distinct particles require a high-rank approximation of *A*. The key idea of our approach is that the difference between the high- and low-rank DMD reconstruction allows us to separate rapidly moving entities from slowly moving or stationary backgrounds. This is first illustrated on simulated kymographs (Fig. S1). To account for anterograde and retrograde movement of motor proteins during IFT, we simulated bi-directional transport in cilia, extracted kymographs and analyzed them by our DMD-based method. Here, the length of the cilium is set to 5 µm corresponding to 80 pixels in our simulation with the stationary protein located close to the tip (Fig. S1A). The IFT trains were blurred to account for the limited resolution of the microscope, and noise was added to create realistic scenarios. Back and forth movement of the bright and dim particles (resembling a fixed number of motors per train) results in a zig-zag movement in the kymograph (Fig. S1B). DMD with TDE and low-rank approximation (here rank, r = 1) captures the background, which is primarily the stationary particle, while DMD with r = 30 allows for precise reconstruction of the simulations (Fig. S1C and D). Key to the success of the DMD approach on our data is the generation of a delay-embedded data matrix, a so-called Hankel matrix, which enriches the kymograph data with time-shifted versions of the simulated transport (see Materials and Methods and below for more details). The eigenvalue spectrum of the decomposed transfer operator is shown in Fig. S1E. By subtracting the low-rank DMD reconstruction from either the original simulation or the high-rank DMD reconstruction, the actively moving component can be isolated, allowing for better discerning the lines of directed transport (Fig. S1F-H). In addition to active transport, ciliary proteins, including motor proteins and certain IFT components, can also enter cilia by diffusion (Ludington et al., 2015). To account for this, we added a diffusing component to the simulation, which resulted in a progressively increasing background component in the simulated video and kymograph over time (Fig. S2A and B). Diffusion was simulated by convolving a constant intensity pool slightly outside of the cilium starting at an arbitrarily chosen start point, ξ (mimicking the ciliary base) with a Gaussian of increasing width. Since the Green’s function of diffusion from a point source is exactly such a Gaussian, this procedure is equivalent to the solution of the 1D-diffusion equation for a fixed concentration corresponding to constant start intensity of a fluorescent entity *u*_0_ at *x* = ξ, (see Supporting Information, Eq. 12) (Socolofsky, 2002). Analyzing such simulations using DMD with TDE reveals not only a high reconstruction quality but also a clear dissection of the complex dynamics into the stationary + diffusing pool (Fig. S2C) and the actively transported component (Fig. S2F and G). These results convincingly demonstrate the unique ability of DMD combined with delay embedding to analyze multiscale transport data from kymographs.

Next, we applied the same DMD-based pre-processing to kymograph data generated from time-lapse recordings of IFT172-eGFP (Fig. 1 and see below). Using a high rank for the SVD step in the DMD-TDE procedure allows for faithful reconstruction of the experimental kymographs (Fig. 1A-C). Using a low-rank approximation (i.e., r = 4, corresponding to only four eigenfunctions/modes) captures the stationary part of the kymograph data (Fig. 1D). Low-rank modes represented stationary or slowly varying background signals, while higher-rank modes captured the fast-moving IFT172-eGFP particles, enabling clear distinction between static and motile populations.

**Figure 1.**
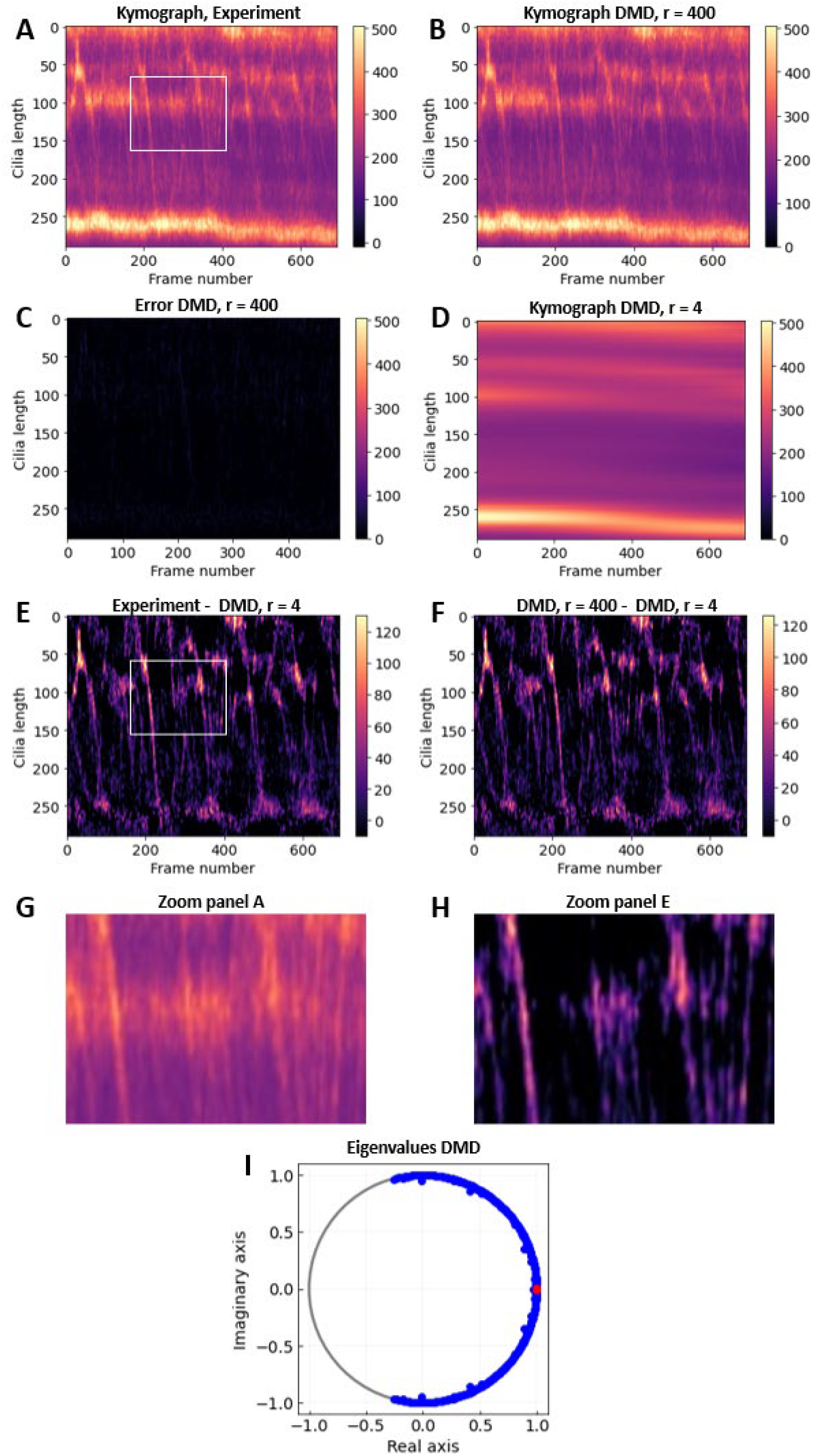
Reconstruction and analysis of experimental kymograph of IFT172-eGFP. A kymograph was generated from a time-lapse video sequence of IFT172-eGFP (A) and reconstructed by DMD with delay embedding dimension, *d* = 200 and rank *r* = 400 (B). The reconstruction error was calculated as L2-norm and shows the high quality of the DMD analysis (C). A low-rank reconstruction with *r* = 4 efficiently isolated the stationary and diffusing pool of IFT172-eGFP in cilia (D). Subtracting this low-rank reconstruction from the experimental data (E) or its high-rank DMD reconstruction (F) reveals the population of IFT172-eGFP moving by active transport visible as straight lines. The white boxes in panels A and E are shown as zoomed versions in G and H, respectively. The eigenvalue spectrum of the reconstructions is also shown (blue dots for *r* = 400 and red dots for *r* = 4 in I).

The low-rank reconstruction of the slow/stationary part can be subtracted from the original data (Fig. 1E) or from the high-rank reconstruction (Fig. 1F) with similar results. This is an efficient method to isolate the protein population moving only by active transport (compare Fig. 1G and H). The eigenvalue spectrum for the low- and high-rank reconstruction is also shown (Fig. 1I). A quantitative comparison of tracks extracted from kymographs demonstrates that pre-processing with DME-TDE indeed increases the length and number of extracted tracks (Fig. S3). Together, our DMD-based isolation of the dynamic protein population facilitates subsequent automated velocity analysis of the pre-processed kymographs.

### A KIF13B mutant does not affect IFT172-eGFP movement in hTERT-RPE1 cells

Given the potential of our method to dissect complex spatiotemporal dynamics of protein transport in cilia, we used it to assess whether KIF13B influences conventional IFT. For that, we used CRISPR-Cas9 methodology to knock out KIF13B in previously characterized hTERT-RPE1 cells stably expressing IFT172-eGFP (Juhl et al., 2023; Kuhns et al., 2024). This, here referred to as “parental”. IFT172, is a component of the IFT-B2 complex (Cole et al., 1998; Pedersen et al., 2005; Taschner et al., 2016) and serves as a reporter for conventional IFT. Two KIF13B mutant clones, B3 and B4, were successfully generated, and their ciliary phenotypes characterized (Fig. 2A; Fig. S4A, D, E, F). Clone B3 had lost the ability to express IFT172-eGFP and displayed significantly reduced ciliary frequency and length compared to parental cells (Fig. S4E, F). Clone B4 expressed IFT172-eGFP, exhibited wild-type ciliary lengths and frequencies (Fig. S4E, F) consistent with previous findings (Rezi et al., 2025; Schou et al., 2017), and was selected for subsequent experiments. This clone expresses a truncated N-terminal fragment of KIF13B (exons 1 to 16) of about 70 kDa, corresponding to the motor, neck-coil (NC), and part of the FHA domain but lacking the C-terminal cargo binding modules (Fig. 2A, B; Fig. S4D). We expect that this truncated KIF13B mutant protein fails to localize to centrosomes and cilia, having lost the RPGRIP1N-C2 type domain that enables binding to NPHP4, required for KIF13B centrosome recruitment (Schou et al., 2017). Similarly, a KIF13B truncation lacking only its CAP-Gly domain has a punctate localization in mouse brain tissue, which is strikingly different from the wild-type protein and comparable to the staining pattern of a full KIF13B knockout line (Mills et al., 2019). Moreover, we expect little to no processive motility on microtubules, as this was shown for different KIF13B proteins truncated within the NC domain or after the FHA domain (Soppina et al., 2014). Cells underwent 24 hours of serum deprivation to induce ciliogenesis, followed by confocal live-cell imaging. We found IFT172-eGFP consistently present inside cilia of both parental and KIF13B mutant cells, where it moved in both anterograde and retrograde directions along the axoneme as expected (Fig. 2C and Movie 1). The velocity of IFT172-eGFP was analyzed from kymographs generated using a MATLAB script that implemented DMD with TDE, effectively decomposing the data into spatial-temporal modes (see above and Materials and Methods for more details; Fig. 2D).

**Figure 2.**
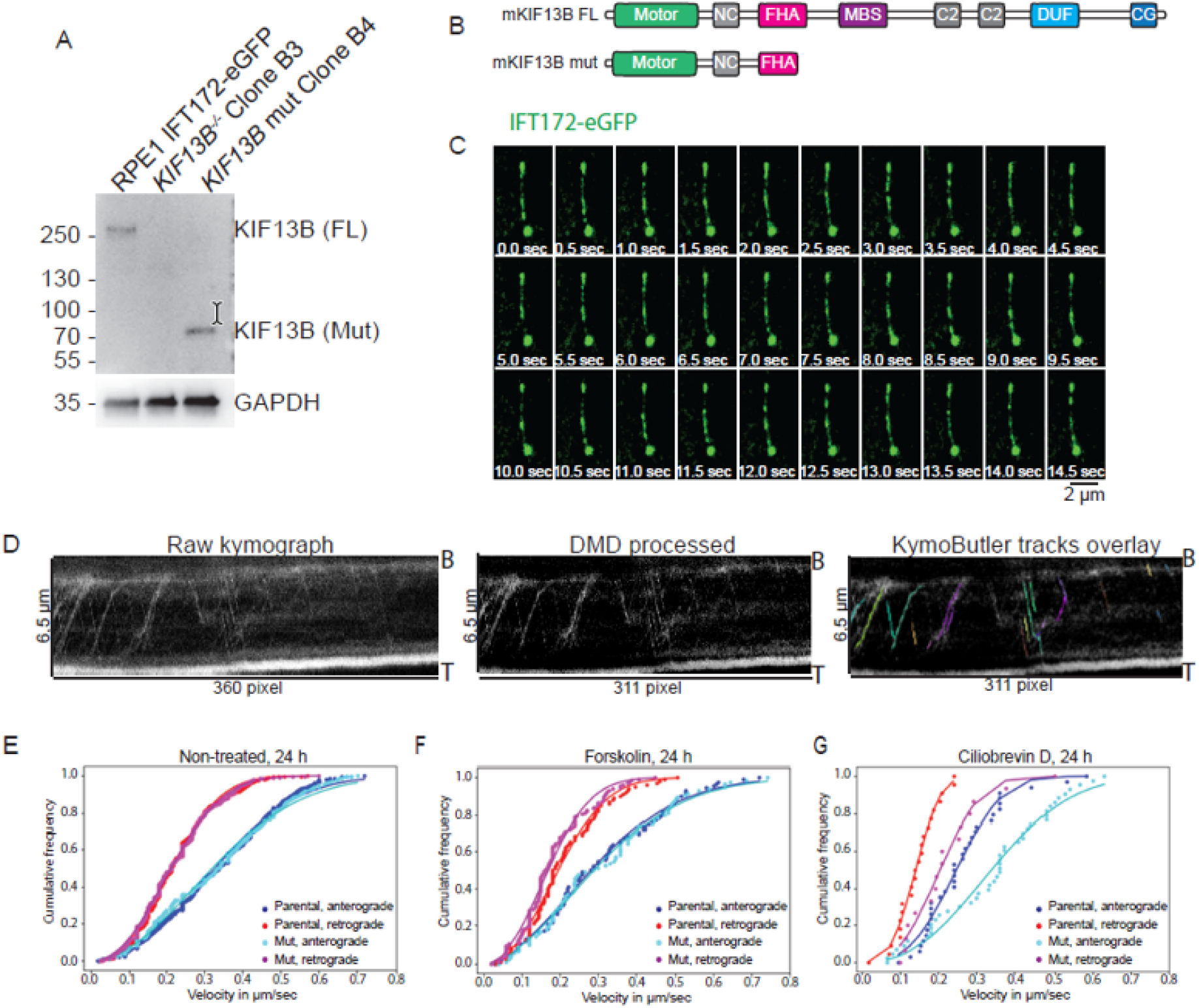
Live-cell imaging of IFT172-eGFP in cilia of 24-hour serum-starved parental and *KIF13B* mutant hTERT-RPE1 cells. (A) Western blot of wild-type (parental) and KIF13B mutant cells (Clone B3, B4) stably expressing IFT172-eGFP. GAPDH: loading control. Molecular mass markers are in kDa. (B) Schematics of protein domains for KIF13B full-length (FL) and clone B4 truncation mutant (Mut). (C) Time-lapse sequence showing IFT172-eGFP (green) moving bidirectionally within the cilium of a parental cell. Based on Movie 1. (D) Representative kymographs depicting cilia of parental cells imaged with high temporal resolution (2 frames/sec) using a confocal microscope with temperature control. b, cilium base; t, cilium tip. From the left: raw kymograph obtained from the MATLAB script; DMD-processed kymograph; overlay of recognized tracks from KymoButler. (E) Cumulative frequency distributions of the velocity of anterograde and retrograde intraciliary movement of IFT172–eGFP in non-treated parental and *KIF13B* mutant (Mut) cells. Graphs are based on 482 (anterograde) and 640 (retrograde) measurements from 32 parental cells, and 335 and 436 measurements from 30 KIF13B mutant cells. (F) Same as (E) but in cells treated with forskolin. Graphs are based on 93 and 102 measurements from 6 parental cells, and 120 and 161 measurements from 7 KIF13B mutant cells. (G) Same as (E) but in cells treated with Ciliobrevin D. Graphs are based on 39 and 24 measurements from 5 parental cells, and 48 and 17 measurements from 5 KIF13B mutant cells. In (E-G), lines show fitted Weibull distribution functions of the form y(v)=1−exp[-(α · v)β].

Subsequent analysis of tracks with the artificial intelligence-based software KymoButler (Jakobs et al., 2019) provided velocities from each kymograph, which were plotted as cumulative distribution function and fitted with a Weibull distribution, as introduced previously (Fig. 2E) (Juhl et al., 2023). This analysis shows that the KIF13B mutant did not affect the shape of the velocity distribution, with anterograde transport of IFT172-eGFP being on average faster than retrograde transport, which can be directly inferred from the Weibull fit parameters (Fig. 2E and Table 1). Indeed, we find that in the parental cells, IFT172-eGFP moves bidirectionally in cilia with an average anterograde velocity of 0.33±0.15 µm/s (mean±SD of n= 482 measurements) and of 0.22±0.10 µm/s in the retrograde direction (mean ± SD of n= 640 measurements from 32 cells) (Fig. 2E). These velocities are in accordance with previous studies (Juhl et al., 2023; Sun et al., 2025), which have shown that anterograde movement is slightly faster than retrograde movement. Similar IFT velocities were observed in KIF13B mutant cells, with average anterograde and retrograde velocities of 0.33±0.16 µm/s (mean±SD of n=335 measurements) and 0.22±0.10 µm/s (mean±SD of n=436 measurements from 30 cells), respectively (Fig. 2E). These results demonstrate that truncation of KIF13B at the C-terminus does not affect IFT as monitored by IFT172-eGFP motility inside cilia under control conditions.

**Table 1.** Mean velocity of anterograde and retrograde IFT172-eGFP, calculated at 24 hours of starvation, for parental and KIF13B mutant cells treated with drugs or vehicle (DMSO) alone. α and β values from Weibull distribution functions are reported.

| Condition | Anterograde |  |  | Retrograde |  |  |
| --- | --- | --- | --- | --- | --- | --- |
| | Mean velocity | $\alpha$ | $\beta$ | Mean velocity | $\alpha$ | $\beta$ |
| Parental, DMSO | $0.33 \pm 0.15 \mu\text{m/s}$ | 2.59 | 2.23 | $0.22 \pm 0.10 \mu\text{m/s}$ | 4.03 | 2.45 |
| Mutant, DMSO | $0.33 \pm 0.16 \mu\text{m/s}$ | 2.60 | 1.99 | $0.22 \pm 0.10 \mu\text{m/s}$ | 4.03 | 2.45 |
| Parental, Ciliobrevin D | $0.26 \pm 0.11 \mu\text{m/s}$ | 3.56 | 2.98 | $0.14 \pm 0.05 \mu\text{m/s}$ | 6.28 | 3.14 |
| Mutant, Ciliobrevin D | $0.34 \pm 0.15 \mu\text{m/s}$ | 2.53 | 2.43 | $0.22 \pm 0.10 \mu\text{m/s}$ | 4.27 | 2.77 |
| Parental, Forskolin | $0.33 \pm 0.14 \mu\text{m/s}$ | 2.89 | 1.85 | $0.19 \pm 0.09 \mu\text{m/s}$ | 4.31 | 2.27 |
| Mutant, Forskolin | $0.31 \pm 0.16 \mu\text{m/s}$ | 2.81 | 1.85 | $0.18 \pm 0.09 \mu\text{m/s}$ | 4.98 | 2.23 |

### KIF13B mutant renders IFT172-eGFP movement insensitive to Ciliobrevin D treatment

To further study whether KIF13B affects IFT, we treated parental and KIF13B mutant cells with the dynein inhibitor Ciliobrevin D, which slows down the velocity of retrograde IFT (Firestone et al., 2012). In parallel, we treated the cells with forskolin, which was reported to accelerate anterograde IFT motility (Besschetnova et al., 2010; Jin et al., 2014). Surprisingly, the treatment with forskolin did not affect the velocity of IFT172-eGFP in either cell line as compared to non-treated cells (Fig. 2E, F), although the cilia appeared to be elongated compared to non-treated cells (Fig. S4I), suggesting that the drug is working. In the case of Ciliobrevin D-treated cells, where cilia length is comparable to that of non-treated cells (Fig. S4I), the retrograde IFT172-eGFP velocity of parental cells is slightly slowed down (0.14±0.05 µm/s, mean±SD of n= 24 measurements) compared to non-treated cells (Fig. 2E, G; Table 1). Interestingly, the KIF13B mutant cells appear to be largely insensitive to Ciliobrevin D treatment, displaying a retrograde IFT172-eGFP velocity of 0.22±0.10 µm/s (mean±SD of n=17 measurements), comparable to non-treated cells (Fig. 2E, G; Table 1). Analysis of stationary IFT172-eGFP particles revealed no significant difference between parental and mutant cells (Fig. S4J, K), suggesting that the insensitivity of KIF13B mutant cells to Ciliobrevin D treatment is not due to altered pausing of IFT trains on the axoneme.

Overall, these results indicate that mutant KIF13B does not drastically affect conventional IFT movement, at least in hTERT-RPE1 cells, but it reduces the sensitity of IFT/dynein-2 to Ciliobrevin D inhibition. It was previously shown that the *C. elegans* KIF13B homologue, KLP-6, slows down the OSM-3 motor and indirectly reduces the velocity of heterotrimeric kinesin-2 in cephalic male cilia (Morsci & Barr, 2011). In our work, we did not observe a significant change in IFT velocity between parental and KIF13B mutant cells, suggesting that KIF13B’s function is distinct from that of KLP-6 although we cannot exclude that the truncated KIF13B mutant protein retains some activity.

### KIF13B and IFT172 move largely independent of each other inside cilia of hTERT-RPE1 cells

To further investigate if KIF13B moves together with IFT trains along the ciliary axoneme, we generated a dual fluorescent reporter cell line stably expressing Halo-KIF13B in the IFT172-eGFP parental background (Fig. S4B, C). The cells were subjected to different starvation times prior to live imaging to assess whether KIF13B shows different activity within the cilia during various stages of cilia formation. Ciliary length and frequency were largely unaffected in the Halo-KIF13B-expressing cells compared to the parental (Fig. S4B, G, H). In coherence with previous findings (Juhl et al., 2023), we observed Halo-KIF13B accumulation at the base of all imaged cilia (n=201) and moving bidirectionally within 22.8% of these cilia (Fig. 3A, B; Table 2; Movie 2;). In some cells, Halo-KIF13B rapidly entered the cilia, remaining up to one minute, and quickly exited from the base (Fig. 3A, B; Movie 2). In rare cases (2.1% of cells starved for 24 hours), we also observed release of vesicle-like particles of Halo-KIF13B from the ciliary tip, followed by rapid exit of KIF13B from the cilium (Movie 3; Table 2 and Fig. S5). We can exclude that dynamic ciliary localization of Halo-KIF13B is due to the fluorescent tag or cilia marker used, as previous work showed the same dynamic behaviour in hTERT-RPE1 cells transiently expressing eGFP-KIF13B in a SMO-tRFP background (Juhl et al., 2023) and in mCCD cells stable expressing mNG-KIF13B with cilia visualized by Sir-Tubulin staining (Rezi et al., 2025). Intriguingly, the percentage of cells with Halo-KIF13B quickly moving into cilia (bursting movement) was higher for short compared to long starvation periods. For example, 29.8% of cilia imaged in cells starved for 6 hours show bidirectional movement of KIF13B inside the cilia, while only 2.5% of the cilia in cells starved for 72 hours present Halo-KIF13B movement (Table 2). Interestingly, movement of IFT172-eGFP was largely unrelated to that of Halo-KIF13B, as these proteins did not co-localize in the traces observed in kymographs (Fig. 3B; Fig. S4). Indeed, when analyzing kymographs of IFT172-eGFP, we often observed diagonal stripes, indicating actively transported proteins and stationary populations (Fig. 1; Fig. 2D; Fig. 3B; Fig S5).

**Table 2:** Overview of imaging results of parental cells expressing Halo-KIF13B, over different starvation times/imaging systems.

| Condition | n | Number of cells with Halo-KIF13B localized to: |  | Comment |
| --- | --- | --- | --- | --- |
|  |  | Ciliary shaft | Tip (release) |  |
| Confocal, Nt, 6 h starvation | 67 | 20 (29.8 %) | / | Live imaging |
| Confocal, Nt, 24 h starvation | 95 | 25 (26.3%) | 2 (2.1%) | Live imaging, two releases from the tip |
| Confocal, Nt, 72 h starvation | 39 | 1 (2.6%) | / | Live imaging |
| SIM, Nt, 24 h starvation | 16 | 5 (31.25%) | / | Fixed cells |

n = total number of cilia imaged; Nt, non-treated.

**Figure 3.**
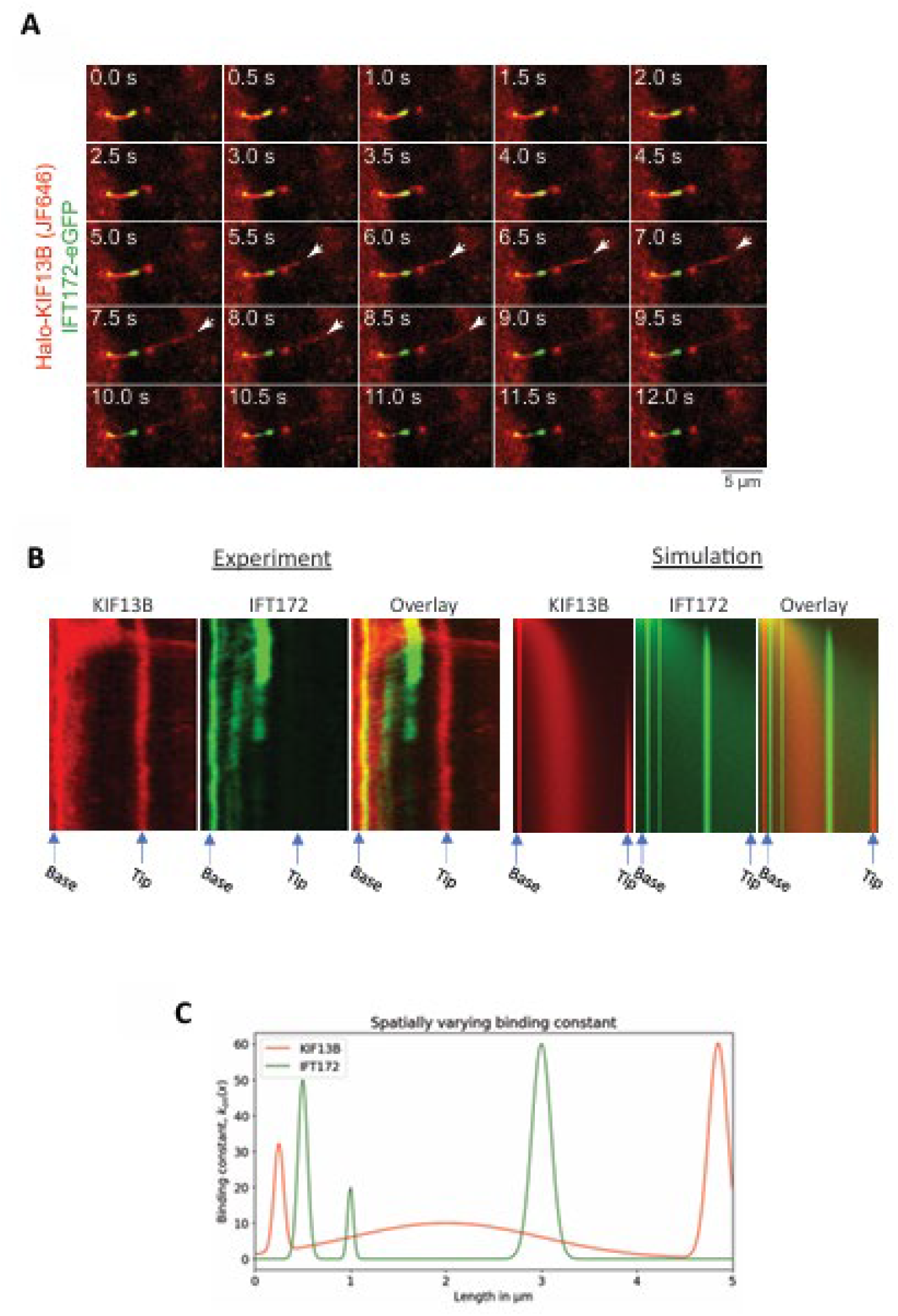
Co-tracking of fluorescent KIF13B and IFT172 reveals distinct motion profiles. Live-cell imaging of cells co-expressing Halo-KIF13B (JF646) (A, red) and IFT172-eGFP (A, green) shows an event of vesicle shedding from the ciliary tip. This vesicle only contains Halo-KIF13B but not IFT172-eGFP. Kymograph analysis from this video sequence additionally reveals distinct transport patterns of both proteins inside the cilium. While a portion of Halo-KIF13B is confined to the base, another pool is spread over the cilium and a third pool is confined to the tip (B, left panels, red). In contrast, IFT172-eGFP is confined in three different regions in the lower part of the cilium (B, left panels, green), not co-inciding with Halo-KIF13B. A numerical simulation of the advection-diffusion equation with a space-dependent binding term can generate kymographs closely matching those observed in experiments (B, right panels). That means, by assuming different binding sites for both proteins inside cilia the spatially differing immobile pools of both proteins can be explained. The binding rate constant *k*_*on*_(*x*) is plotted for both simulated protein populations (C).

### Advection-diffusion and transient binding underlie the complex dynamics of proteins in cilia

For Halo-KIF13B the dominating signal in the kymographs consisted of a broadly distributed pool and one or several immobile populations (Fig. 3B, left panel). The broadly distributed pool is likely a combination of freely diffusing and broadly bound KIF13B proteins. Furthermore, the confined immobile populations observed for both Halo-KIF13B and IFT172-eGFP (horizontal stripes) were found at different locations inside cilia (Fig. 3B). In some kymographs, we also found a diffusive component apparently moving along with the active populations, particularly for Halo-KIF13B (Fig. 3B; Fig. S5). Mathematically, active transport (so-called advection) gives rise to the diagonal stripe-like pattern, observed for both proteins but being more dominant for IFT172-eGFP. Combined with diffusion this can be described by an advection-diffusion equation with velocity *v* and diffusion coefficient *D*, respectively (see Materials and Methods for details). Such a model arises naturally from a molecular description of kinesin motors and other proteins moving along filaments if the observed transport times are long compared to the attachment and detachment rates (Smith & Simmons, 2001). While active transport translocates either Halo-KIF13B or IFT172-eGFP along the cilium, diffusion causes spreading of its components. Both scenarios are illustrated in Fig. S6A-D and can be faithfully reconstructed using DMD with TDE (Fig. S6E, F). As further explained in the Appendix, the ability of DMD-TDE to reconstruct advection-diffusion based transport in cilia is a consequence of the fact, that the Koopman-based time-shift operator, used in delay embedding, corresponds to first order to the spatial drift term used in the advection-diffusion equation (see Fig. S7 for an illustration).

We can show in additional simulations, that combining advection-diffusion of proteins with binding at distinct sites inside cilia can visually reproduce the observed kymographs for Halo-KIF13B in the IFT172-eGFP, respectively (Fig. 3B, C). In particular, the presence of a diffusion barrier can also be simulated, revealing a sharpe edge at the site of the barrier, in contrast to the stripe-like pattern found for a membrane-bound protein population (Fig. S8). While such edge-like patterns were not observed in the experimental kymographs, we cannot rule out that diffusion barriers contribute to the regulated access of both proteins to cilia. Also, additional simulations reveal that the unavoidable photobleaching of some of the moving protein populations results in an underestimation of the contribution of diffusion to IFT of fluorescent proteins (Fig. S9). Together, our combined experimental and computational analysis suggests that both KIF13B and IFT172 show distinct motion patterns inside cilia due to differences in active transport, transient binding events and eventual barriers to diffusion.

### KIF13B can be rapidly released from the membrane and exported from cilia

To better understand the differences in motion pattern of KIF13B and IFT172, we carried out additional time-lapse experiments, focusing on a cell with a slightly thicker cilium, in which the lumen can be clearly resolved by confocal time-lapse imaging (Fig. 4A, B). Here, we found that Halo-KIF13B localizes to the membrane, while IFT172-eGFP is localized in the center of the cilium. KIF13B suddenly leaves the cilium while IFT172-eGFP is still visible within, suggesting that the cilium stays in focus. The integrated intensity of Halo-KIF13B (red signal in Fig. 4A) was measured in the cilium as outlined from the IFT172-eGFP signal upon dilating the binary mask by two pixels (Fig. 4B). Quantification of Halo-KIF13B’ s intensity in this mask reveals a sudden intensity drop followed by a stable plateau showing that the membrane-associated protein pool leaves the cilium and becomes confined to the base or pocket, i.e. underneath the green IFT172-eGFP spot at the ciliary base (Fig. 4B, C). This is also clearly seen from the kymographs (Fig. 4D-F). Since the membrane signal of Halo-KIF13B gets averaged when measured in this way (because of averaging the signal across the thickness of the cilium in a kymograph), the line profile across the cilium close to the base (Fig. 4G) was additionally plotted for selected time points in Fig. 4H. This makes the membrane localization of Halo-KIF13B clearer (Fig. 4H, red lines) and underlines that it is segregated from the luminally located IFT172-eGFP (Fig. 4H, green lines). We speculated that the exit of Halo-KIF13B from the cilium is too fast to be happening by diffusion alone. Therefore, the advection-diffusion equation for an open domain on one end and reflecting boundary at the other was solved and the function fitted to the data, starting at the beginning of the drop in cilia intensity (red box in Fig. 4C; Fig. 4I). The fit described the data very well and provides an estimate for the diffusion coefficient of Halo-KIF13B of D = 0.59 µm^2^/s and for the speed of active transport of v = 0.92 µm/s. Notably, this velocity is slightly higher than the maximal retrograde velocity measured from kymographs for Halo-KIF13B moving inside the cilia (compare Fig. 2E and Fig. 4I). After export of Halo-KIF13B, only a fluorescent pool at the base, slightly underneath that of IFT172-eGFP, remains, giving rise to a stable fluorescence plateau (compare Fig. 4B and C). The background intensity B = 75.64 resembles this pool of Halo-KIF13B confined to the cilia base plus eventual remaining luminal fluorescence, which is slightly higher than the intensity of the exported fraction, i.e. amplitude was fixed to 55 intensity units (not shown). Thus, we propose that Halo-KIF13B binds to membrane zones within cilia, from which it is suddenly released by a combination of diffusion and directed transport (advection).

**Figure 4.**
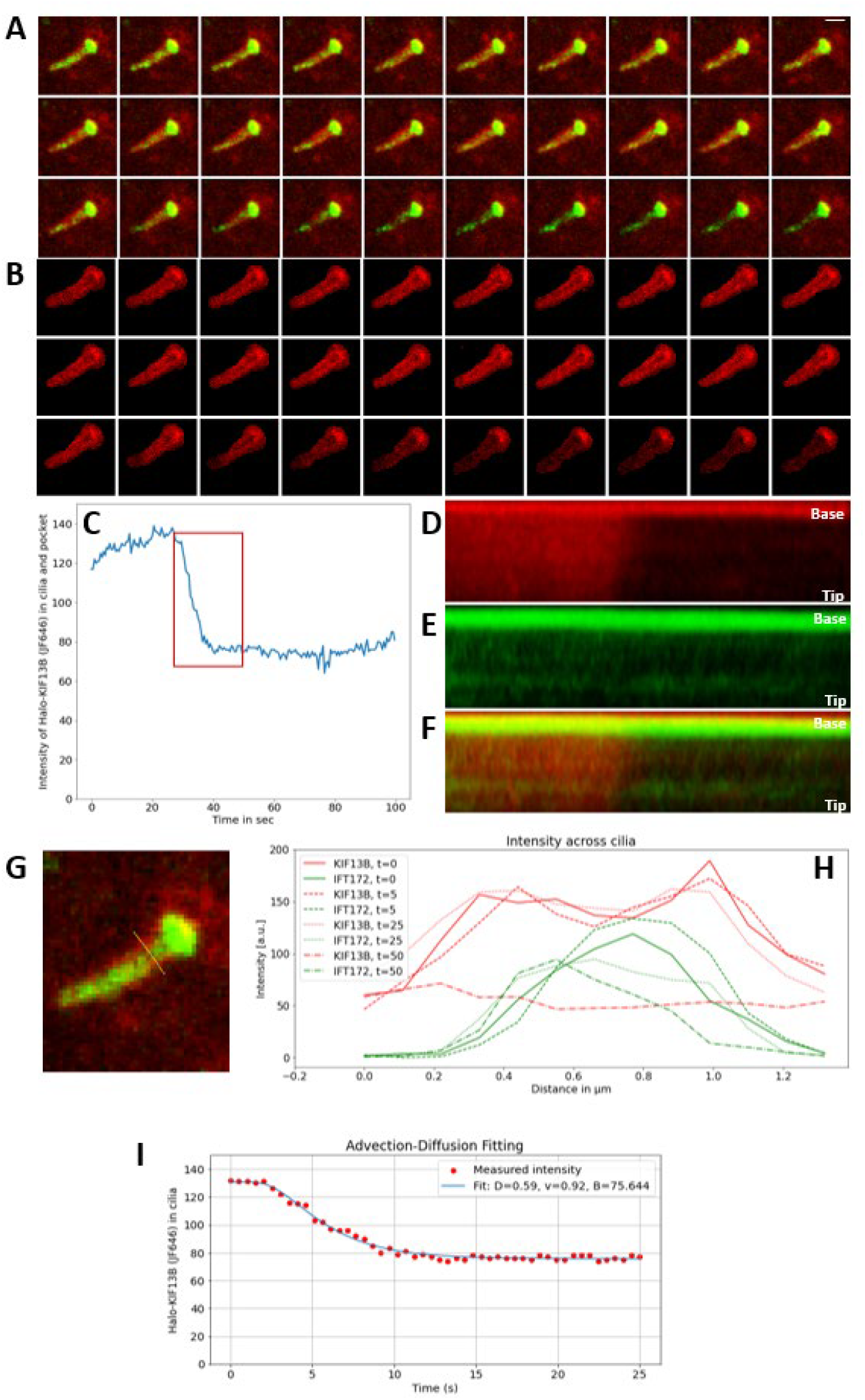
Quantiative time-lapse analysis reveals transient membrane association of KIF13B. Live-cell imaging of cells expressing both markers on a confocal microscope revealed membrane localization of Halo-KIF13B (JF646) (A, red) but not of IFT172-eGFP (A, green), the latter being confined to the cilia lumen and base. Based on the intensity mask for IFT172-eGFP dilated by two pixels, the fluorescence of Halo-KIF13B (JF646) is shown, revealing sudden membrane release and export from the cilium (B). Quantification of this signal supports rapid export of labeled KIF13B from the cilium (C). Kymographs of Halo-KIF13B (JF646) (D), IFT172-eGFP (E) and color overlay (F) are shown. Zoomed version with line scan across cilium close to the base (G) from which intensity profiles where extracted at selected time points (H). An advection diffusion model with reflecting boundary conditions on one site and absorbing boundary on the opposite site to model release of protein from cilia was fitted to the time-dependent integrated intensity of Halo-KIF13B (JF646) (I, compare red box in C). The model fits the data well and provides a diffusion constant of D = 0.59 µm^2^/s and a transport velocity of v = 0.92 µm/s.

### KIF13B localizes close to the ciliary and periciliary membrane compartments

To further explore KIF13B behavior and localization in cilia, we performed Structured Illumination Microscopy (SIM) on fixed cells. We observed Halo-KIF13B accumulation at the ciliary base, as for live-cell imaging, where it appeared to accumulate at two distinct sites: one below IFT172-eGFP fluorescence and one overlapping with IFT172-eGFP at the base (Fig. 5A, B). Moreover, in 31.2% of the cilia visualized with this method (n = 16), we could see Halo-KIF13B localized in the cilia shaft (Table 2). SIM imaging revealed that Halo-KIF13B fluorescence is only rarely co-localized with IFT172-eGFP across the axoneme, and the two proteins mostly retain different distributions (Fig. 5B). IFT172-eGFP is more internally localized than Halo-KIF13B in the cilia shaft, exhibiting a discrete punctate pattern spanning across 0.2 µm of the cilia diameter (Fig. 5B, C). IFT172 can indeed associate with the internal region of the axoneme and its fibrous core in *Trypanosoma brucei* (Alves et al., 2024), although it was also reported to interact with membranes (Wang et al., 2018). In the IFT172-eGFP- and Halo-KIF13B-expressing cells, KIF13B appears to localize external to IFT172-eGFP, showing two distinct fluorescent peaks of 0.1 µm each, spanning across an “empty”, non-fluorescent region of 0.1 µm, where IFT172-eGFP is located (Fig. 5B, C). This ciliary localization pattern of Halo-KIF13B is similar to that of the membrane-spanning receptor Htr6, as visualized by Stimulated Emission Depletion microscopy (STED) (Kohli et al., 2017). These results suggest that KIF13B resides in proximity to the ciliary membrane, where it may play a scaffolding role to promote endocytic retrieval (Rezi et al., 2025), similar to its proposed scaffolding during recruitment and endocytosis of LRP1 in hepatocytes (Kanai et al., 2014). Since KIF13B is also often localized at the base of the cilia, this localization was investigated using STED imaging of fixed IFT172-eGFP and Halo-KIF13B expressing cells using a FBF-1 antibody to visualize the basal body transition fibers/distal appensages (Fig. 5D). Here IFT172-eGFP is found within the cilium and FBF-1 is seen as the classical ring structure (Kazatskaya et al., 2017), where Halo-KIF13B is located just beneath it in both mother and daughter centrioles.

**Figure 5.**
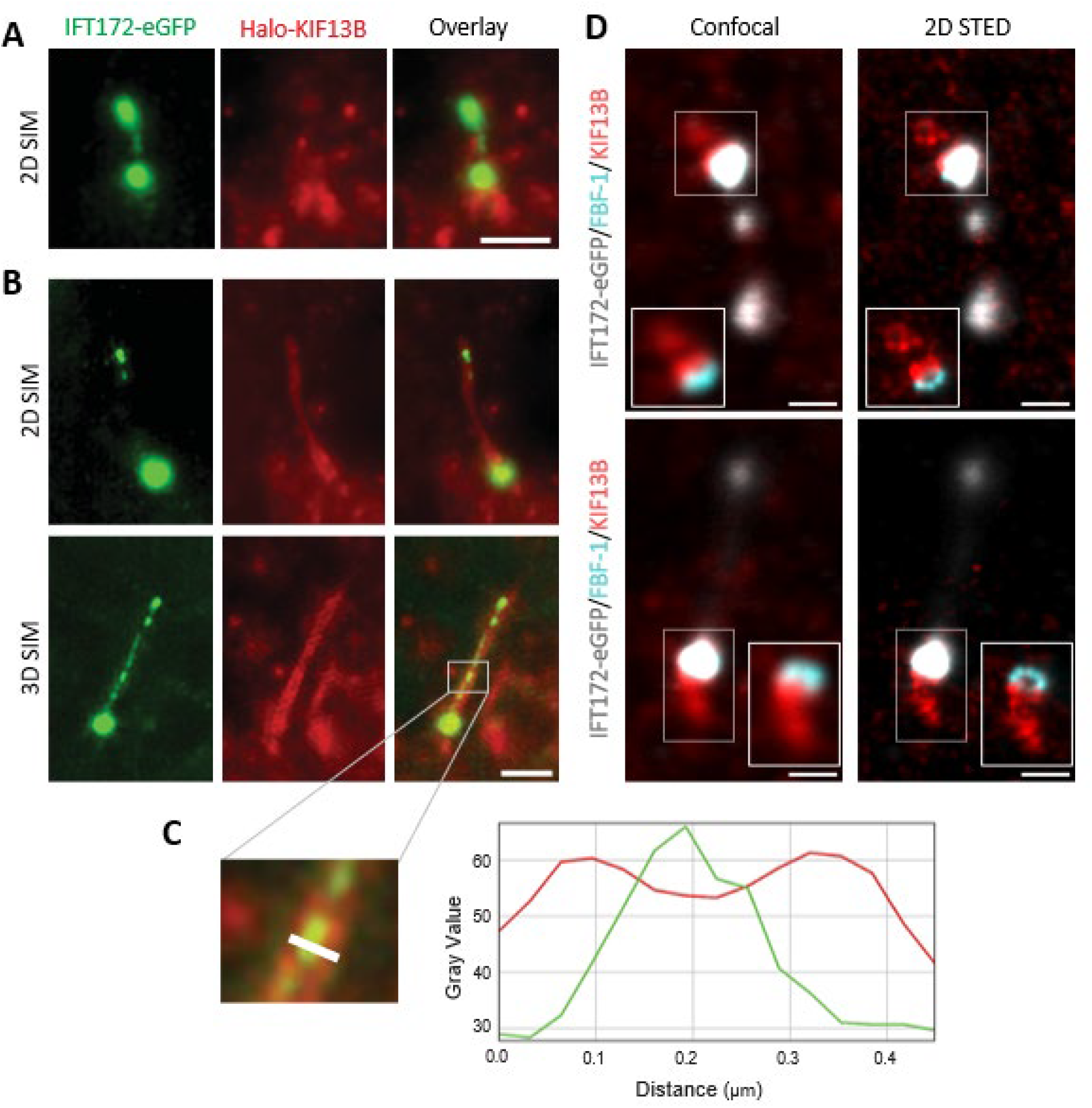
Super-resolution imaging reveals membrane localization of Halo-KIF13B. (A, B) 2D (A) and 3D (B) SIM imaging of Halo-KIF13B (red) and IFT172-eGFP (green) stably expressed in hTERT-RPE1 cells subjected to 24 hours of starvation. In panel B, the white line was used to calculate fluorescence intensity profiles. Scale bars are 2 µm. (C) Fluorescence intensity profile calculated for the same cilium as in (B), drawing a line near the ciliary base, perpendicular to the axoneme. (D) Confocal (right) and 2D STED (left) images of fixed hTERT-RPE1 cells expressing Halo-KIF13B (red) and IFT172-eGFP (gray) after 24 hours of starvation, with FBF-1 antibody staining (cyan). Here, a Gaussian blur with a sigma of 2 pixels is used to smooth the images. Scale bars are 1 µm.

## Conclusions

Our work combines quantitative live-cell imaging with computationally improved kymograph analysis as well as simulation and regression of partial differential equations to discern protein transport in primary cilia. Using these tools, we demonstrate that the kinesin motor KIF13B is transported in cilia with a combination of active and diffusive transport but largely independent of classical IFT. On the other hand, removing most of the cargo-binding tail region of KIF13B does not affect IFT172/IFT-B2 velocities within cilia, indicating that KIF13B’s action within cilia is unlikely to rely on physical association with the IFT machinery. However, the KIF13B truncating mutation seems to render the cells less sensitive to Ciliobrevin D treatment. A study in *C. elegans* showed that loss of the transition zone (TZ) protein NPHP4, which interacts physically with KIF13B (Schou et al., 2017), rescues the ciliary exit and velocity of retrograde IFT trains in *wdr-60* null mutants, which display an underpowered dynein-2 motor due to loss of the intermediate chain subunit WDR60 (De-Castro et al., 2022). This is in accordance with the known gating function of the TZ (Nachury, 2018; Park & Leroux, 2022; Ye et al., 2018). The interaction with NPHP4 therefore places KIF13B in a functional relationship with the TZ barrier and dynein-2-dependent retrograde transport. Surprisingly, however, STED super-resolution imaging presented here reveals that KIF13B localizes not at the TZ but rather at a more proximal region coinciding with the subdistal appendages (sDAs) of the mother centriole. The ability of KIF13B to modulate IFT sensitivity to dynein-2 inhibition must therefore operate independently of TZ barrier function and instead implicates the sDAs as a previously unrecognized regulatory node for IFT dynamics. This interpretation is supported by a growing body of evidence linking sDA-associated proteins to dynein-2 and kinesin regulation at the ciliary base. CEP170, a core sDA component, has recently been shown to interact directly with dynein-2, promoting complex assembly at the sDAs (Weijman et al., 2024). This raises the possibility that the sDA platform coordinates the reciprocal regulation of dynein-2 and kinesin-2/kinesin-3 motors prior to their entry into the ciliary compartment. In this model, KIF13B positioning near the sDAs and at the cytoplasmic microtubule anchoring point would be strategically suited to two functions: first, enabling efficient trafficking of the kinesin into the cilium during bursts, and second, facilitating KIF13B-mediated regulation of endocytic vesicle trafficking away from the ciliary base, consistent with its previously known role in vesicle modulation and interactions with vesicle-associated proteins (Cho et al., 2023; Morthorst et al., 2018; Rezi et al., 2025). This dual role of the sDAs as a scaffold for both motor complex assembly and periciliary vesicle trafficking is further supported by our previous finding that loss of the sDA protein CEP128 in hTERT-RPE1 cells impairs RAB11 recruitment to the ciliary base and disrupts TGF-β/BMP-dependent phosphorylation of kinesin-3 family members KIF1B, KIF1C, and KIF13A (Mönnich et al., 2018). RAB11-positive recycling endosomes are well-established carriers of ciliary cargo and signaling receptors, and their impaired recruitment in CEP128 knockout cells suggests that the sDA scaffold is required for the local endosomal organization at the ciliary base. The fact that KIF13B also localizes to this compartment and interacts with vesicle-associated proteins places it within the same regulatory axis.

We show by SIM analysis that KIF13B is co-localized with the ciliary and periciliary membrane compartment, when moving inside cilia in bidirectional bursts. So far the only other protein known to exhibit similar behavior is the Rac/Rab-effector MiniBAR, which also co-localizes with the ciliary membrane and moves bidirectionally inside the ciliary shaft (Serres et al., 2023). Moreover, the number of cilia that show MiniBAR ciliary bursts decreases progressively during the advancement of ciliogenesis, similarly to KIF13B. Actin polymerization is necessary for the reorganization of the ciliary tip compartment and consequent ectocytosis (Kiesel et al., 2020; Loukil et al., 2021; Nager et al., 2017; Phua et al., 2017; Prasai et al.), while local clearance of F-actin at the ciliary base is necessary for cilia elongation (Brücker et al., 2020). MiniBAR is shown to limit actin-dependent contractility by inhibiting Rho GTPase activity, promoting ciliogenesis (Serres et al., 2023). It has been demonstrated that KIF13B negatively regulates ectosome release in cilia (Rezi et al., 2025), and it localizes to the hepatocyte side of the actin-rich microvilli in the liver (Kanai et al., 2014). Moreover, KIF13B directly binds to Centaurin-α1 (ADAP1) (Tong et al., 2010), suppressing its ARF6 GAP activity, which indirectly leads to elevated ARF6-GTP levels that can influence actin polymerization and dynamics (Venkateswarlu et al., 2005). When KIF13B encounters phosphoinositides like PIP3 or PI(3,4)P2, the release of ADAP1 is triggered, restoring its GAP function (Duellberg et al., 2021). It would therefore be interesting to explore whether KIF13B interacts functionally with MiniBAR or independently exerts a similar role of actin regulation inside the cilia shaft and/or the ciliary tip. Another aspect to examine further would be the involvement of KIF13B in membrane curvature sensing, as KIF13B significantly impairs the wedge-driven curvature exosome biogenesis pathway (Rezi et al., 2025), interacts with the BAR domain-containing Angiomotin p80 (Morthorst et al., 2022), and BAR domains can promote vesicular budding, sensing, and contribute to membrane curvature (Habermann, 2004). Finally, it remains to be determined exactly which signal triggers the ciliary entrance and burst-like movement of KIF13B, and further work is also needed to explore the mechanisms involved. One possibility is that KIF13B associates with intraluminal vesicles, which have been proposed to carry some cargo into cilia, for example G-protein coupled receptors (Tingey et al., 2024). Transient binding to actively moving vesicles inside cilia could also contribute to the advection-diffusion type motion for KIF13B and IFT172 (Wang et al., 2018), but further work is needed to clarify the identity and role of such vesicles in transport inside the primary cilium.

## MATERIALS AND METHODS

### Reagents and plasmids

An overview of reagents and plasmids described in this study is shown in Table S1.

To create transgenic cell lines for live-cell imaging, we generated a lentiviral expression construct coding for Halo-KIF13B, expressed under IRES and EF1a promoters to ensure stable and low expression (Aslanyan et al., 2023). Shortly, the full-length human *KIF13B* coding sequence was cloned from pENTR20-mNG-KIF13B-FL into pENTR20-Halo-C1 by digesting both plasmids with NotI and KpnI restriction enzymes followed by standard ligation. The resulting Gateway entry plasmids were further recombined with the pCDH-EF1a-Gateway-IRES-BLAST destination plasmid using the Gateway LR recombination kit (Invitrogen, Cat#11791020). All cloning vectors were kindly supplied by Dr. Kay Oliver Schink, Oslo University Hospital, Norway, and were described in (Campeau et al., 2009). *Escherichia coli* DH10α was used for transformation according to standard procedures. Plasmid purification was performed using the NucleoBond Xtra Midi EF Kit from Macherey-Nagel (Cat# 740420.50).

To generate lentiviruses, lentiviral expression plasmids were co-transfected with packaging plasmids pMD2.G and pCMVΔ-R8.2 (kind gift of Dr. Carlo Rivolta, Institute of Molecular and Clinical Ophthalmology Basel, Switzerland) into HEK293T cells using Lipofectamine 3000 (Invitrogen, Cat#L3000015) according to the manufacturer’s instructions. The clarified culture medium containing the lentivirus particles was subsequently collected.

### Mammalian cell culture and stable cell lines generation

HEK293T cells used in the study are laboratory stocks purchased from ATCC (cat# CRL-3216). The hTERT-RPE1 parental cell line stably expressing IFT172-eGFP (derived from the immortalized hTERT-RPE1 cell line, ATCC, clone CRL-4000) has been described previously (Juhl et al., 2023; Kuhns et al., 2024). hTERT-RPE1 stably co-expressing IFT172-eGFP and Halo-KIF13B were generated by transducing the aforementioned cell line with lentiviral particles expressing pCDH-EF1aGW-IRES-Blast-Halo-KIF13B plasmid. Pools of parental cells stably expressing Halo-KIF13B were selected with 10 µg/ml Blasticidin S HCL (Gibco, Cat#r210-01). After 7 days of selection, cells expressing the reporter gene were further selected based on the Janelia Fluor 646 Halo ligand fluorescence, with a FACS Aria III instrument at the FACS Facility at Biotech Research & Innovation Centre (University of Copenhagen, Copenhagen, DK). Successful expression was confirmed through western blotting and IFM. Parental cells expressing mutant KIF13B were generated by CRISPR-Cas9. Briefly, SpCas9 2NLS Nuclease (Synthego) and CRISPRevolution sgRNA EZ Kit containing two sgRNA against the beginning of *Kif13b* exon 17, purchased from Synthego database (Table S1), were co-transfected into the parental cells using the Nucleofector with P3 Primary Cell 4D-Nucleofector® X Kit S (Lonza Bioscience Cat# V4XP-3032) following manufacturer’s instructions. Program EA-104 was selected on the 4D Nucleofector System Core Unit (Lonza Bioscience, Cat# AAF-1003B), and the cuvettes containing the cells were placed in the X Unit (Lonza Bioscience, Cat#AAF-1003X). Cells were subsequently replated in single-cell dilution in a 96-well plate by manual limiting dilution. After reaching confluency, the cells were subcultured into two 96-well plates: one for screening the clones for KIF13B depletion by western blot analysis, and the other for maintaining the clones in culture. Analysis led to the selection of two clones with *KIF13B* mutations (Clone B3 and B4), B4 was used in this study. PCR-amplified genomic DNA from the genomic area surrounding the Cas9 cut site of the selected clones was sent to Eurofins Genomics for Sanger sequencing. It revealed several deletions in the intronic area before *KIF13B* exon 17 for both clones (Fig. S1C). Cell lines were routinely checked for contamination.

Cells were grown in an incubator at 37°C, with 5% CO2 in Dulbecco’s modified Eagle’s high-glucose medium (DMEM, Gibco, Cat#41966-052) supplemented with 10% heat-inactivated fetal bovine serum (FBS, Gibco, Cat #10438-026) and 1% penicillin-streptomycin (Sigma-Aldrich, Cat#P0781). Cell cultures were passaged every 3–4 days, unless stated otherwise. To induce ciliogenesis, cells were cultured for 24 h in serum-starvation medium (DMEM free of FBS and antibiotics). For Ciliobrevin D and forskolin experiments, culture medium was exchanged with fresh medium containing 10 µM Ciliobrevin D (Merck, Cat# 250401) or 100 µM forskolin (Merck, Cat# F6886). Imaging started after 3 h, and control cells treated with a similar volume of DMSO were analyzed in parallel.

### SDS-PAGE and western blot analysis

SDS-PAGE and western blotting were performed according to the procedures described by (Gonçalves et al., 2021). Antibodies and dilutions employed are listed in Table S1. Original immunoblots are shown in Fig. S2. Briefly, cells were lysed with M-PER lysis buffer (ThermoFisher, Cat# 78501), transferred to Eppendorf tubes, and incubated for 5 minutes on ice. After centrifugation at 20,000 x g for 15 min at 4 °C to remove debris, the supernatant was collected. Protein concentrations were determined using BioDrop µLITE (Biochrom, Cat# 4AJ-6319282). Lysates were prepared for SDS-PAGE analysis by adding NuPAGE™ LDS Sample Buffer (4X) from Thermo Fischer Scientific (Cat#NP0007) and 50 mM DTT, followed by 5 minutes heating at 95 ºC. SDS-PAGE was performed with the Mini-PROTEAN® Tetra System and Mini-PROTEAN® TGX^TM^ Precast Gel 4-15% 10 wells from Bio-Rab Laboratories, Inc. (Cat#456-1083). PageRuler^TM^ Plus Prestained Protein Ladder (Thermo Fischer Scientific, Cat#26619) was used as molecular mass marker. The Trans-BLOT® Turbo Transfer System from Bio-Rad Laboratories, Inc. was used for the transfer of proteins from gels onto Trans-Blot® Turbo^TM^ Transfer Pack, Mini format 0.2 μm Nitrocellulose membranes (Bio-Rad Laboratories, Inc.; Cat#1704158). Membranes were blocked in 5% bovine serum albumin (BSA) in Tris-buffered saline with Tween-20 (TBS-T; 10 mM Tris-HCl, pH 7.5, 150 mM NaCl, 0.1% Tween-20) for 1 hour at room temperature and then incubated for 1,5 hours at room temperature or overnight at 4 ºC with primary antibodies diluted in 5% BSA in TBS-T. Membranes were washed three times for 10 min at room temperature in TBS-T prior to incubation with secondary antibodies diluted in 5% BSA in TBS-T for 45 minutes at room temperature. Finally, after three 10-minute washes at room temperature in TBS-T, blots were developed with SuperSignal^TM^ West Pico PLUS chemiluminescent Substrate (Thermo Fischer Scientific, Cat#34580) and visualized using a FUSION FX SPECTRA machine from Vilber.

### Immunofluorescence microscopy

For IFM analysis, cells were seeded on glass coverslips (confocal imaging) or on µ-slide 8 well with #1.5H glass coverslip bottom (cat. no. 80807, ibidi GmbH) (SIM and STED imaging) to obtain 80% confluency and subjected to 24 hours of serum starvation before fixation. Cells were washed in ice-cold PBS, fixed with 4% PFA for 15 min at room temperature (confocal imaging) or at 4 °C (SIM and STED imaging), and cell membranes were permeabilized by incubation with 1%(w/v) bovine serum albumin (BSA) and 0.2% (v/v) TritonX-100 in 1× PBS for 12 min at room temperature. To avoid non-specific binding of antibodies, a 1-hour blocking step at room temperature with 2% BSA in 1× PBS was performed, followed by incubation with primary antibodies for 1.5 hours at room temperature or overnight at 4°C. After three 5-minutes washing steps with 2% BSA in 1× PBS, cells were incubated for 45 min at room temperature, with the appropriate Alexa Fluor-conjugated secondary antibodies diluted in 2% BSA in 1× PBS, and nuclei were labeled with DAPI (Sigma-Aldrich, Cat# D9542). Coverslips were mounted with Shandon Immu-Mount (Thermo Fisher Scientific, Cat# 9990402) supplemented with 0.5% N-propyl gallate onto glass slides. For SIM and STED imaging, no antibodies were used to visualize Halo-KIF13B and IFT172-eGFP, as they showed Janelia Fluor ligand-mediated and intrinsic fluorescence, respectively.

Confocal fluorescence images were captured on a motorized and automated Olympus IX83 inverted microscope, equipped with a Yokogawa CSU-W1 confocal scanner unit, Prime 95B back-illuminated sCMOS camera (Teledyne Photometrics), and Olympus UPlanSApo 100x oil microscope objective. We used the Olympus CellSens Dimension software, version 4.2 (Evident). Images for publication were processed with FIJI (Schindelin et al., 2012), Adobe Photoshop version 26.4.1 (Adobe), and Adobe Illustrator version 29.1 (Adobe). Structured Illumination Microscopy and 3D SIM Imaging were performed on a Nikon N-SIM build onto a Ti-2 microscope body, with an SR HP Apo TIRF 100xAC oil objective for imaging. STED imaging was performed on an Abberior Facility Line STED microscope (Abberior Instruments GmbH) using a 60x NA 1.4 objective (UPLXAPO60XO). All excitation and STED lasers are pulsed and circularly polarized. IFT172-eGFP was excited with 488 nm, and collected on a matrix detector between 498-551 nm. STAR ORANGE was excited with 561 nm, and collected on an APD between 580-630 nm. Halo-KIF13B labelled with Janelia Fluor 646 was excited with 640 nm, and collected on an APD between 650-763 nm. For STED imaging, both STAR ORANGE and Janelia Fluor 646 were depleted using 775 nm.

### Live-cell fluorescence imaging

To observe the dynamics of IFT172-eGFP and Halo-KIF13B, the cells were cultured on 35 mm glass-bottom dishes (WillCo Wells, Cat# HBST-3522) and serum-starved for 24 hours. Janelia Fluor 646 (Promega, Cat# GA1120) was used as a fluorescent ligand for Halo-KIF13B. Cells were incubated in culture starvation medium with 0.05 µM of Janelia Fluor 646 for 10 minutes, followed by three washes in PBS. As a primary cilia marker, SiR-tubulin (Cytoskeleton, Inc., Cat#CY-SC002) or Spy555-tubulin (Cytoskeleton, Inc., Cat# CY-SC203) staining was used. The starvation medium was supplemented with 1 µM of SiR-tubulin or 1:1000 Spy555-tubulin, and the cells were incubated at 37 °C and 5% CO2 for at least 30 min prior to cell imaging. For the treatment with Ciliobrevin D and forskolin, the staining with fluorescent probes was done during the last 30 min of drug incubation. All live-cell imaging was carried out in DMEM or M1 medium. Imaging was performed on an Olympus inverted microscope (IX83) equipped with a Yokogawa spinning disc and on a Nikon Ti-2, A1 confocal microscope with a Plan Apo λ 100× NA 1.4 oil objective (at DaMBIC, University of Southern Denmark). Both microscopes were equipped with a humidity chamber, enabling temperature regulation (37°C) and CO2 (5%) control. The following laser settings were used: 488 nm for eGFP, emission measured between 500-550 nm, 561 nm for Spy555-tubulin emission measured between 570-620 nm, and 640 nm for Janelia Fluor646 and SiR-Tubulin-Cy5 emission measured between 663-738 nm. The interval for the time-lapse sequences was set to 0.5 s/frame.

### Background on Dynamic Mode Decomposition (DMD) and kymograph analysis

For DMD analysis of live-cell imaging data, the kymographs are seen as space-time matrices, in which each column is a snapshot in time (i.e., the intensity measurement along the line from which the kymograph was generated. In addition to the kymograph matrix starting at the first time point (Eq. 1, below), a time-shifted version starting at the second time point (Eq. 2, below) is generated leading to a sequence of snapshots arranged into two matrices (Schmid, 2010; Tirunagari et al., 2019):

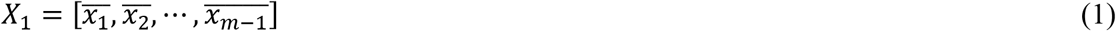

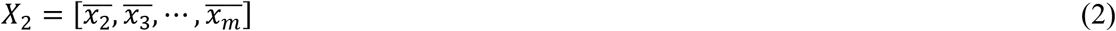

Each line of a kymograph is one snapshot, and represents a column vector of pixel positions along the line, 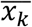. The index *k*=1,…, *m* runs over all acquired snapshots in time with steps, Δt. One can define the progression from state x(k·Δt)=x_k_ to x_k+1_ as:

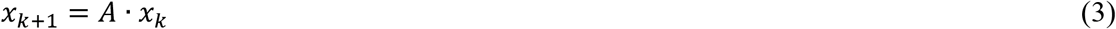

Here, *A* is a matrix which describes the advancement of the system from image *x*_k_ to image *x*_k+1_. This matrix resembles the Koopman or transfer operator for measurements g(*x*_k_)=*x*_k_ and the goal of DMD is to find a rank-truncated approximation to the matrix *A* (Brunton & Kutz, 2019). A known limitation of DMD is its inability to reconstruct purely translational movement (advection) accurately, and to overcome this limitation, various modifications have been developed, including multiresolution DMD and Lagrangian DMD (Kutz et al., 2015; Wu et al., 2021). The latter, was designed for analysis of advection-dominant processes by DMD using a particle-centered coordinate system, thereby outperforming standard DMD. Here, we show that standard DMD can be used for the analysis of advection-dominant processes, such as active transport in cilia, when time-delay embedding (TDE) is applied before the analysis. TDE uses time-sifted versions of the original data stacked into a large, so-called Hankel matrix, thereby enriching the original spatiotemporal sequences with past information (Pan & Duraisamy, 2020). For implement TDE, one must enrich the data matrices of Eqs. 1 and 2 with past observations, thereby efficiently increasing the spectral content of the data. This is realized by constructing a Hankel matrix of the data matrix of Eq. 1 by including time-shifted versions of each row according to:

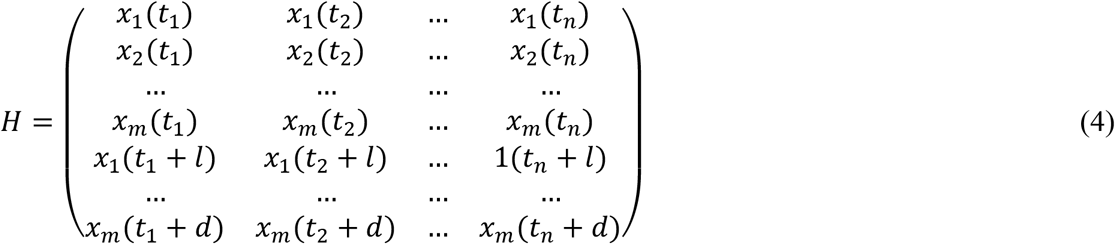

Here, *t*_i_, *i* = 1, …*n* are the snapshots and *x*_j_ = 1, …*m* are the individual time traces, i.e., the simulated or experimental time courses for each cell. The index *l* is the time delay (chosen as *l* = 1 or 5) and *d* is the embedding dimension, i.e., the total number of chosen time shifts. The matrix H and its time-advanced variant were used in the standard DMD algorithm starting from Eqs. 1 to 3 and consisting of a rank-truncated SVD and minimization of the Frobenius norm followed by spectral decomposition of the resulting matrix *A*_*r*_, as described in previous publications (Brunton & Kutz, 2019; Wustner et al., 2024; Wustner et al., 2025).

### Additional image analysis and statistics

Image data from IFM were processed and analysed quantitatively using methods previously described in (Rezi et al., 2025). The measurements of cilia length and frequency in parental, fluorescent reporter, and mutant cell lines were done using FIJI software (Schindelin et al., 2012) and were normalized against the parental cells control, and subsequent statistical analysis was performed using GraphPad Prism 10 software. All quantitative data are presented as mean ± standard deviation (SD). Statistical significance was determined using an unpaired ANOVA test or an unpaired t-test, and defined as a p-value < 0.05. Unless otherwise noted, all the experiments were repeated in at least three independent biological replicates. Significance levels are indicated as followed: p > 0.05 not significant (ns), p ≤ 0.05 *, p ≤ 0.01, ** p ≤ 0.001 ***, p ≤ 0.0001, ****.

Movies from live-cell imaging were processed with a self-developed script in MATLAB (available from the authors upon request). The segmented cilia were first spatially aligned using a rigid-body registration procedure. A line (straight or segmented) was drawn along the cilia length to select the cilia shaft, from where the kymographs were extracted, and then they were refined with DMD with time-delay embedding implementation for improved resolution as described in Appendix 1. Kymograph tracks were recognized using the deep learning software KymoButler (Jakobs et al., 2019) in Wolfram. The UniKymoButler method was used, with the following parameters to recognize single unidirectional tracks: threshold = 0.2, minimum frames = 3, minimum object size = 3. Velocities were determined based on the known pixel size and time interval. Cumulative histograms of velocities measured in the anterograde and retrograde directions were calculated and fitted to a Weibull function of the form:

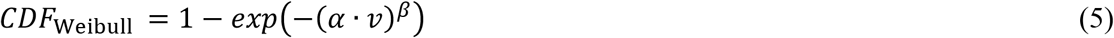

Here, α is the rate constant of how fast the CDF curve changes as function of velocities, *v*, β is the stretching parameter and *CDF* _Weibull_ is the cumulative distribution function.

### Analytical solution of the advection-diffusion equation and the effect of photobleaching

The advection-diffusion equation reads for the one-dimensional case:

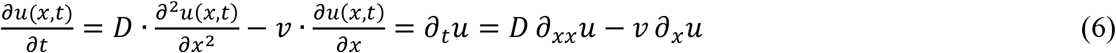

Here, *u*(*x,t*) is the fluorescence intensity of the moving species, which in our case is a protein moving actively by motor-driven transport with velocity *v* inside cilia. On the right side, we use short-hand notation for the time-derivative ∂/∂*t* = ∂_*t*_, as well as for the first and the second spatial derivative, i.e., ∂/∂*x* = ∂_*x*_ and ∂^2^/∂*x*^2^ = ∂_*xx*_, respectively. In addition to active transport modeled by the drift term, *v ∂*_*x*_*u*, proteins can be released from the IFT trains and diffuse with diffusion constant *D* in the cilia space. This diffusion process is here assumed to be one-dimensional and corresponds to the term *C ∂*_*xx*_*u*. For an initial Gaussian concentration profile located at the left border (*x* = 0), the solution to Eq. 6 can be found analytically e.g., by Fourier transformation in space, solving the corresponding ordinary differential equation and transforming back (not shown). This leads for an initial δ-injection of the mass *M* at the cilia base and Dirichlet boundary conditions to (Socolofsky, 2002):

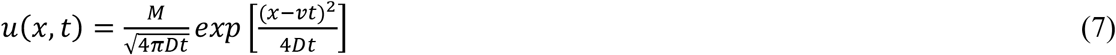

In case of Neuman boundary conditions (i.e. reflection at the boundaries, *L*), the solution can be found with the method of images and reads for a constant protein supply from the ciliary base (Socolofsky, 2002):

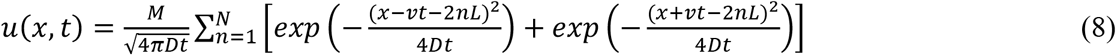

In the presence of homogeneous photobleaching in cilia with rate constant *k*, this expression becomes (Socolofsky, 2002):

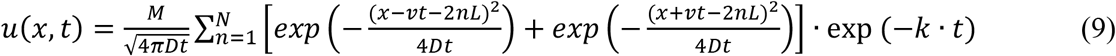

This solution was plotted for varying diffusion constants, *D*, velocities, *v*, and bleach rate constants, *k*, as described in the main text.

### Numerical simulation of advection-diffusion and advection-diffusion-reaction equations

To simulate the solution of the advection-diffusion equation for a spatially varying diffusion constant, D(x), the spatial coordinate, *x*, was discretized into 400 grid points along the entire cilium length of L = 5 µm. The time step, dt, was chosen to be very small to avoid numerical instabilities in the presence of advection (dt = 2·10^-4^ sec), ensuring that *v*·d*t*/d*x* ≤ 1. An initial Gaussian profile was simulated at the base, x = 0.08 µm and width of 0.03 µm, approximating a sharp peak of protein intensity entering the cilium by diffusion and advection (optional). Diffusion was implemented as finite-volume and advection as explicit upwind scheme (Kuzmin, 2010). Space-dependent diffusion was implemented as Gaussian-shaped dip at *x* = 1.5 µm and width (sigma) of 0.15 µm from *D* = 0.5 µm^2^/sec to 0.05 µm^2^/sec. To implement the space-dependent diffusion process numerically, Eq.6 was rewritten as:

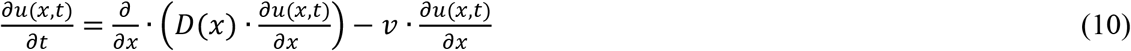

A Crank-Nicholson scheme was chosen for discretization of the diffusion term (Kuzmin, 2010). At the boundaries, von Neumann conditions were implemented. To include spatially varying binding events, the advection-diffusion-reaction has to be simulated, which reads:

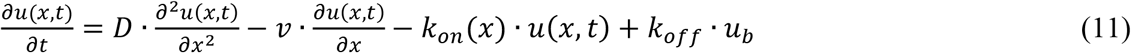

Here, the last two terms on the right-hand site described space-dependent binding with rate constant *k*_*on*_(*x*) and dissociation with (spatially fixed) off-rate constant, *k*_*off*_, respectively. The spatial profile of *k*_*on*_(*x*) was varied between the two studied proteins and is shown in Fig. 3C. A reflecting boundary was chosen at the top of the cilium, while an absorbing boundary at the bottom ensures that protein can be transported back into the cell body during retrograde transport.

### Fitting of advection-diffusion-binding model to export kinetics of Halo-KIF13B from cilia

The analytical solution of Eq. 7 was implemented in a Python code using the Scipy library (Virtanen et al., 2020) upon including a background term, B, to carry out a fit to the experimental data of Halo-KIF13B export from cilia (see Results section, Fig. 4). Reflecting boundary at the base and absorbing boundary at the tip were included, and the diffusion constant, *D*, the background term, *B*, and the transport velocity, *v*, were estimated from the data.

### Stochastic simulation of bi-directional protein motion inside cilia to validate DMD-TDE

Random (Brownian) motion combined with a drift term switching direction at the top and base of cilia was implemented in ImageJ (https://imagej.net/ij/) as described previously (Juhl et al., 2023). To emulate a stationary population a Brownian particle moving very slowly (i.e. in a region approximating the particle size) was added close to the tip of the cilum in the simulation. Next, this simulation was extended by including a freely diffusing protein population, which is constantly released at the ciliary base and diffuses into the cilium with diffusion constant, *D*. The solution of the 1D-diffusion equation for a fixed protein concentration corresponding to a constant intensity *u*_0_ at *x* = ξ < 0, reads (Socolofsky, 2002):

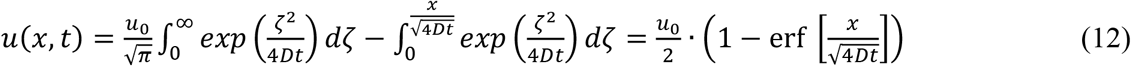

Here, 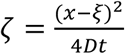 and ‘erf’ is the error function. This solution was added to the stochastic simulation output shown in Fig. S1 and plotted as function of space coordinate x and time, *t*, producing the simulation output shown in Fig. S2.

## Appendix

### Time delay embedding approximates the advection-diffusion operator

Time-delay embedding (TDE) was originally introduced to reconstruct the attractor of a dynamical system from measured time series by embedding it in a higher-dimensional space (Takens, 1980). More recently, it has been shown to be effective in constructing finite-dimensional approximations of nonlinear dynamics within the Koopman operator framework (Kamb et al., 2020; Pan & Duraisamy, 2020). Delay coordinates are defined with time delay τ and embedding dimension *d* as:

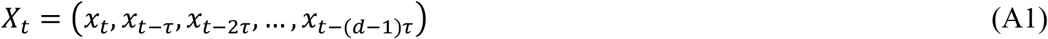

Each time delay corresponds to the application of the inverse time-shift operator T^-*k*^ with *k* = τ shifting the time series *backward* in time by τ, which leads to:

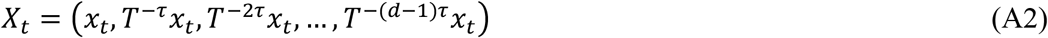

Thus, each delay corresponds to the application of a backward time-shift operator. For τ = Δt, the operator shifts the time series backward in discrete steps, as used in the Hankel matrix, see Eq. 4:

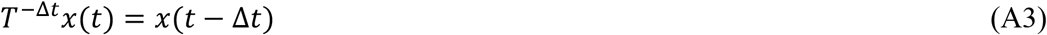

For a smooth signal, we can expand the right-hand side into a Taylor series:

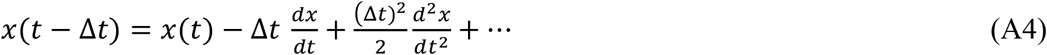

This corresponds to the series expansion of the exponential operator:

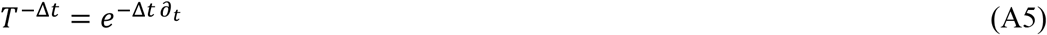

Applying this to a spatially distributed observable *u*(*x, t*) and truncating at first order, one gets:

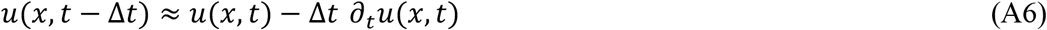

Using the advection–diffusion equation (see Eq. 6 in the main text) we obtain:

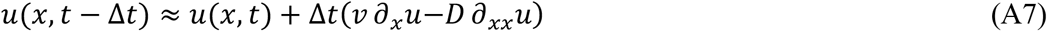

A Taylor expansion in space gives:

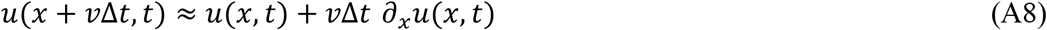

Combining both expressions, Eq. A7 and A8, yields:

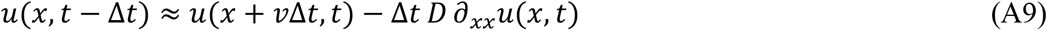

This shows that a backward shift in time corresponds, to first order, to a forward shift in space, corrected by diffusion.

Defining a spatial shift operator *S*^+1^by *S*^+1^*u*(*x, t*) = *u*(*x* ^+^ Δ*x, t*), we obtain:

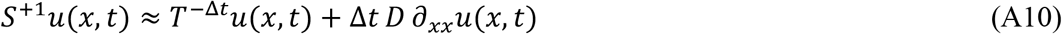

Thus, temporal shifts encoded in delay coordinates correspond to spatial translations governed by the underlying dynamics. This explains why TDE provides DMD with access to the relevant Koopman-invariant subspace (Das & Giannakis, 2019). The observable *uu*(*xx, tt*), i.e. the measured fluorescence intensity, evolves under the Koopman operator, whose generator is the advection– diffusion operator. By incorporating delayed snapshots, DMD can approximate this evolution more accurately, leading to improved reconstruction of advective transport. Delay embedding can thereby be understood as a first-order approximation of the Koopman time-shift operator associated with advection–diffusion dynamics.

## Acknowledgements

We thank Dr. Kay Oliver Schink for the plasmids and Søren Johansen for technical assistance. We thank the Danish Molecular Biomedical Imaging Center, University of Southern Denmark, for the use of imaging equipment. We thank Drs. Stefanie Kuhns and Jens S. Andersen, University of Southern Denmark, for an aliquot of the FBF-1 primary antibody and for hTERT-RPE1 cells stably expressing IFT172-eGFP. We thank Rajesh Somasundaram at the Flowcytometry facility at BRIC - Biotech Research & Innovation Centre, University of Copenhagen, for help with FACS sorting of fluorescent reporter cell lines.

## Author contributions

Conceptualization: L.B.P., D.W.; Methodology: L.B.P., D.W, F.C., L.L.; Validation: L.B.P., D.W, F.C., L.L.; Formal analysis: F.C. L.L., D.W.; Investigation: L.B.P., D.W, F.C., L.L.; Model development: DW; Writing-original draft: F.C.; Writing-review & editing: F.C., L.L., L.B.P, D.W.; Visualization: L.B.P., D.W, F.C., L.L.; Supervision: L.B.P., D.W.; Project administration: L.B.P.; Funding acquisition: L.B.P., D.W.

