## Supplemental material for "Analysis of motor-based transport in primary cilia by dynamic mode decomposition of live-cell imaging data"

q

[2]Department of Biochemistry and Molecular Biology, University of Southern Denmark, Campusvej 55, DK-5230 Odense M, Denmark

### Supporting figures

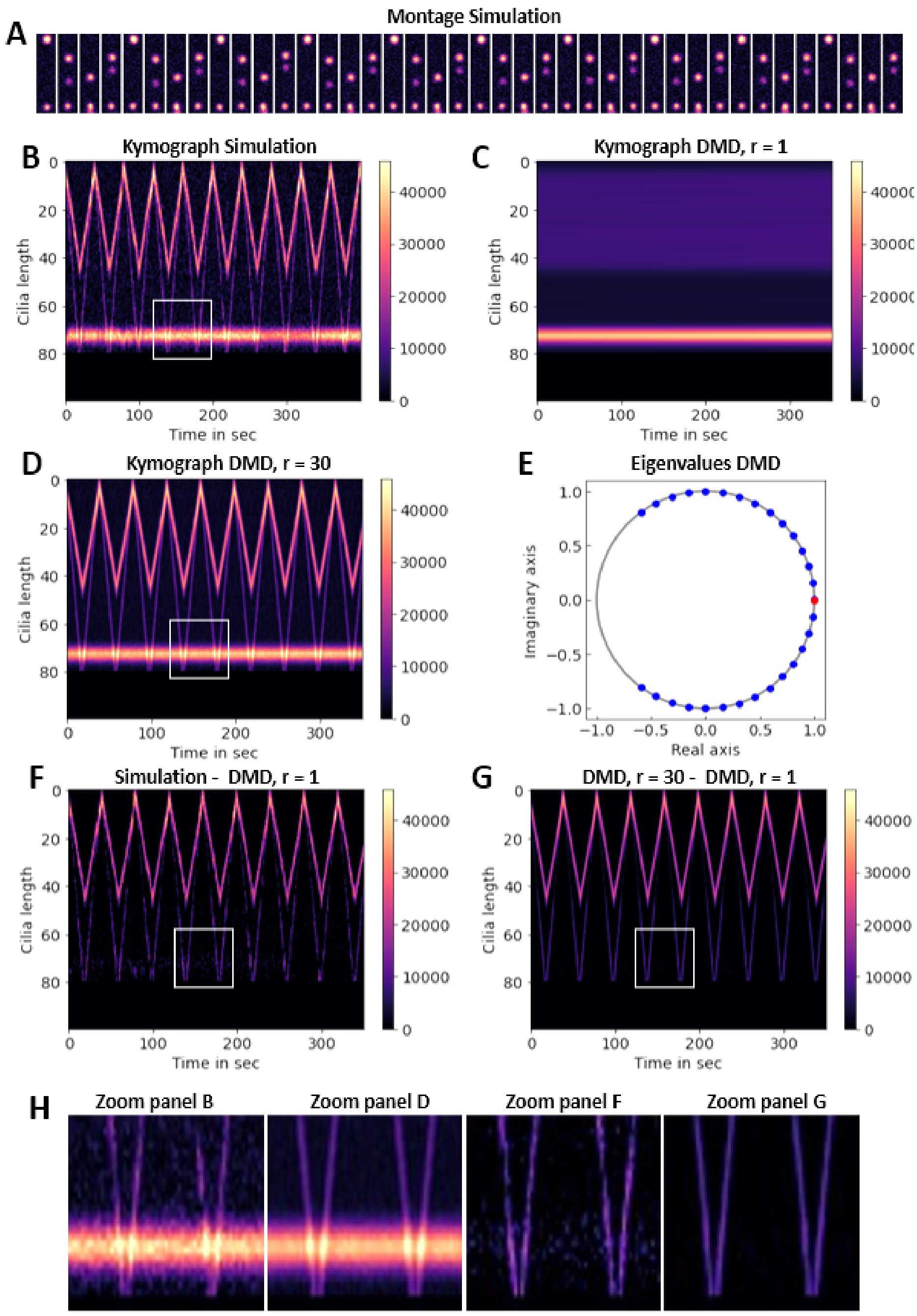


**Figure S1. Simulation and reconstruction of motor-based transport by DMD with delay embedding.** Motor movement in both directions along cilia was simulated for two IFT trains with differing velocities and a third population, which remained stationary, only showing slight oscillations around a mean position at the cilia tip (bottom in A). Simulation shapshots (A) and kymograph of the data (B) were generated in ImageJ. Reconstruction by DMD after delay embedding with a delay, d = 50 was carried out for rank r = 1 (C) or r = 30 (D). Spectral decomposition of the system matrix, A, provides eigenvalues for both reconstructions (blue dots for r = 30 and red dots for r = 1 in E). Subtraction of low-rank DMD reconstruction from simulation (F) and from high-rank reconstruction (G). Zoomed regions indicated by white boxes in B, D, F and G are enlarged, revealing the efficient removal of the stationary protein pool (H).

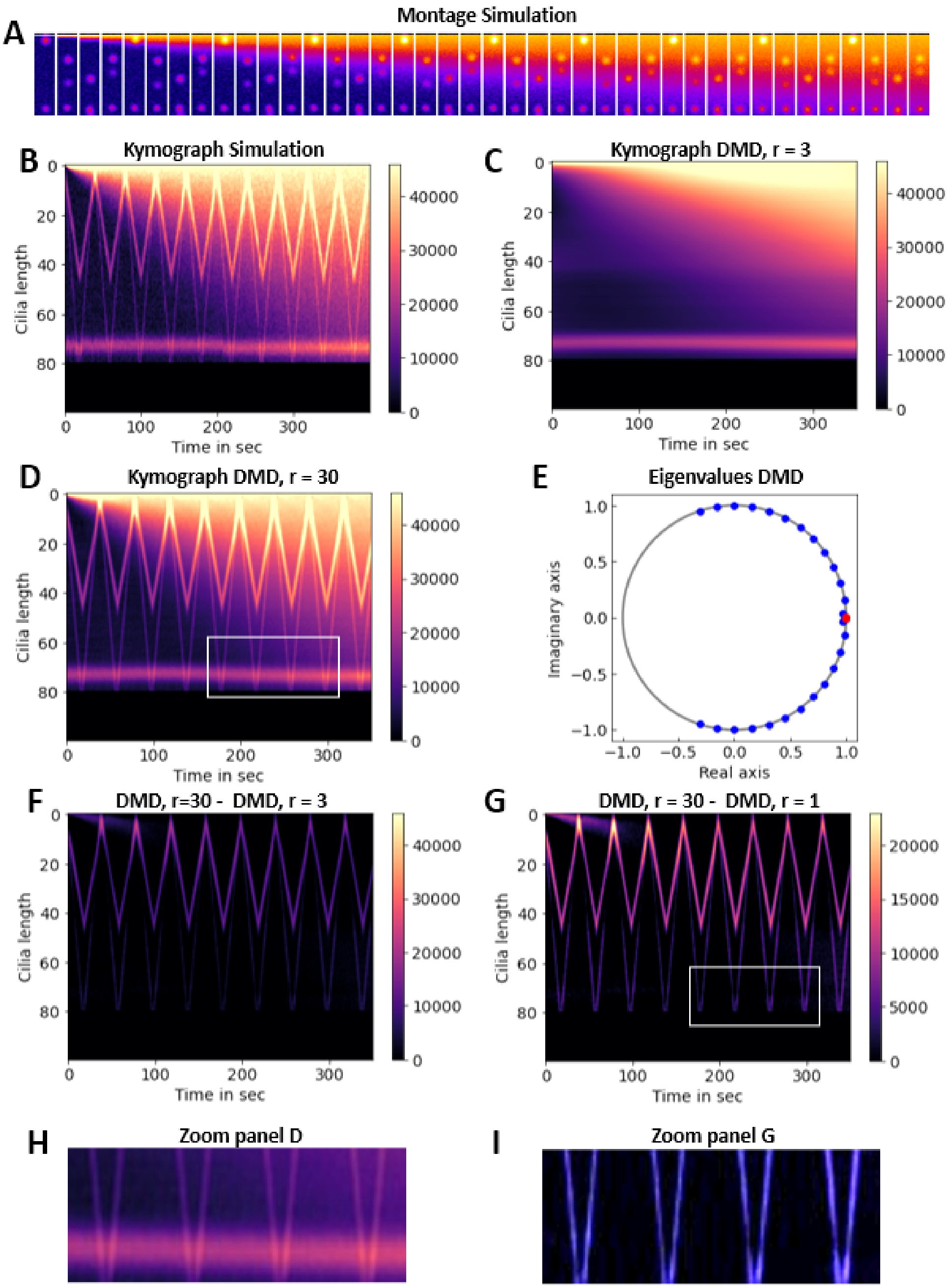


**Figure S2. DMD reconstruction of simulated motor transport with diffusing species.** Bi-directional motor movement was simulated along cilia for two IFT trains with differing velocities and a third population, which is stationary, only showing slight oscillations around a mean position at the cilia tip (bottom ). A population was added, which only moves by diffusion from the base into the cilium (top in A). Snapshots of the simulation (A) and kymograph of the data (B) were generated in ImageJ. DMD-TDE reconstruction after delay embedding with $d = 50$ was carried out for rank $r$ =3 (C) and subtracted from the simulated data (D). Spectral decomposition of the system matrix, A, provides eigenvalues for DMD reconstructions (blue dots for $r = 30$ and red dots for $r = 3$ in E). Zoomed regions indicated by white boxes in B and D are enlarged, revealing the combination of a pool moving by directed transport, diffusion and a stationary pool in the simulated data (F) and the efficient removal of the stationary and the diffusing protein pools by the DMD procedure (G).

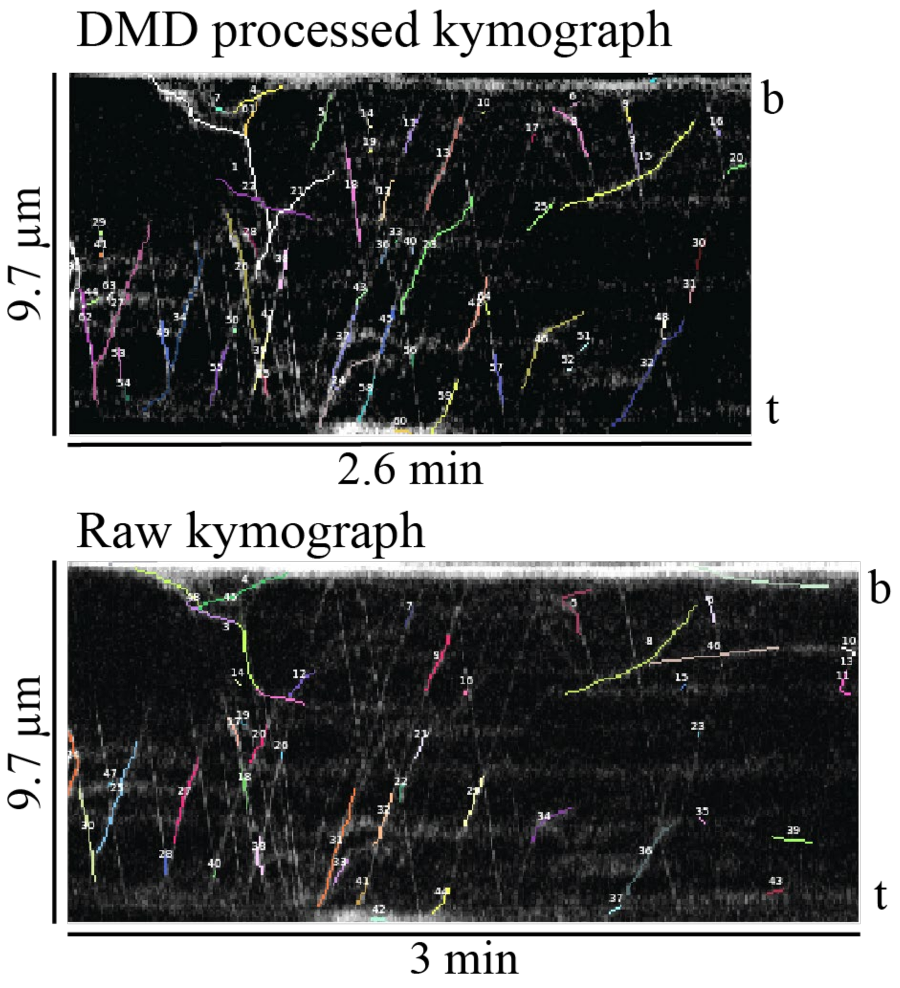


**Figure S3. DMD pre-processing improves kymograph analysis.** Using DMD with TDE as described in Materials and methods provides more and longer tracks (upper panel) compared to unprocessed kymographs (lower panel). Due to the delay-embedding the kymograph is shortened, though, in the DMD-TDE case. Nevertheless, our method enables KymoButler to detect 65 traces in the pre-processed kymographs but only 49 in the raw kymographs.

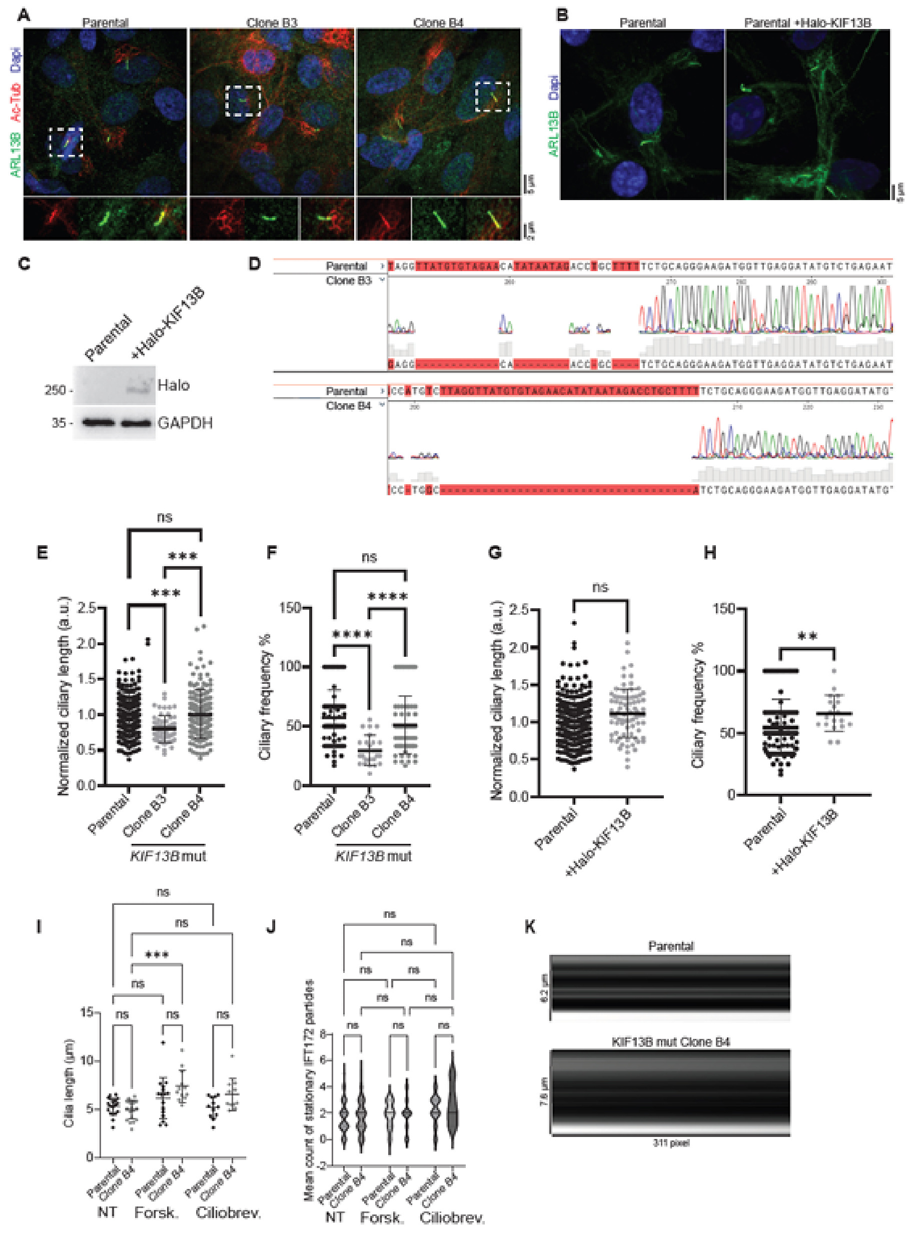


**Figure S4. Analysis of stable and drug-treated hTERT RPE1 cell lines.** (A) IFM of IFT172-eGFP-expressing parental and KIF13B mutant cells. Acetylated α-tubulin (AcTub) and ARL13B mark the cilium. Insets show enlargement of the cilium-centrosome axis. (B) IFM of parental cells with or without stable expression of Halo-KIF13B. ARL13B marks the cilium. Insets show enlargement of the cilium-centrosome axis. (C) Western blot of parental- and Halo-KIF13B cells. GAPDH: this is the loading control. Molecular mass markers are in kDa. (D) Sanger sequencing of the KIF13B mutant clones B3 and B4. B3 shows multiple deletions in the intronic region before exon 17, B4 shows a 36 bp deletion in the same region. (E-H) Quantification of ciliary length (E, G) and frequency (F, H) of indicated cell lines. Ciliary length was normalized to the parental line. Graphs show data as mean ± SD, with significance determined using an unpaired ANOVA test. (I) Quantification of cilia length in cells treated with DMSO alone (NT), Ciliobrevin-D (Ciliobrev.), or forskolin (Forsk.). (J) Mean count of stationary particles of IFT172-eGFP in NT cells, treated with Ciliobrevin-D, or treated with forskolin. (K) Kymographs with isolated stationary movement of IFT172-eGFP, obtained by the dynamic mode decomposition (DMD) with time-delay embedding.

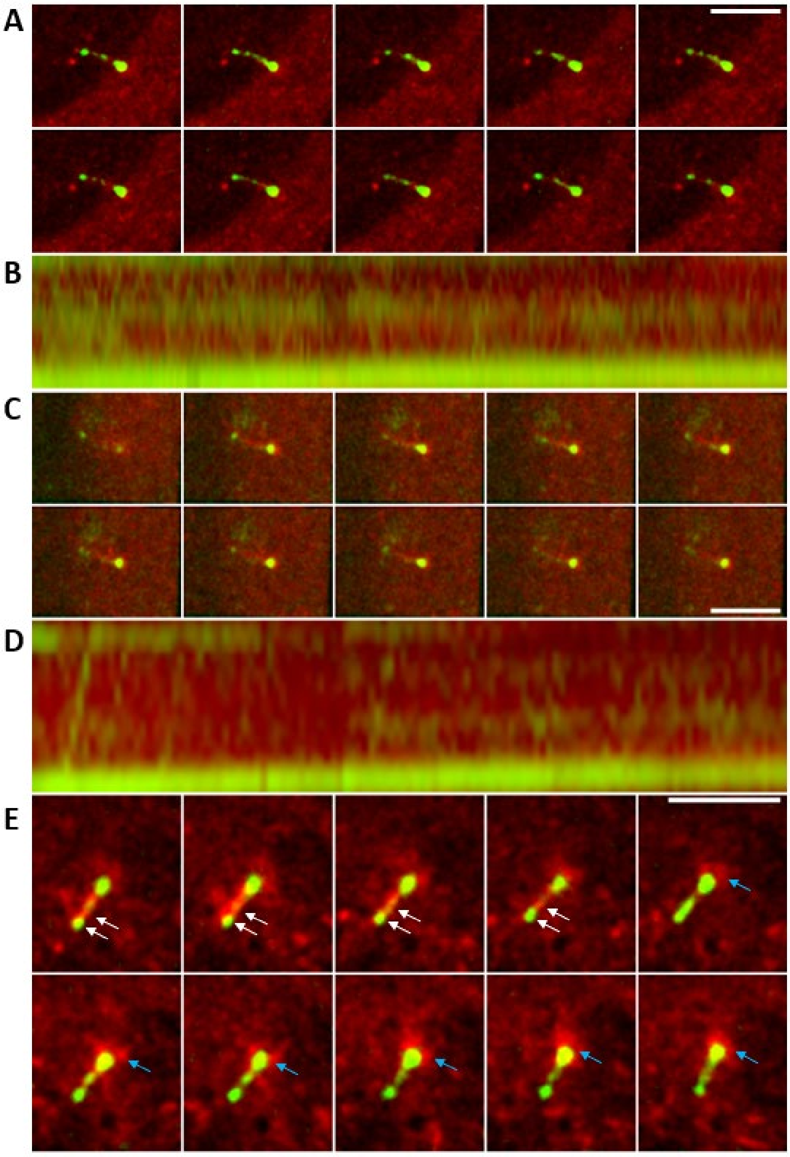


**Figure S5. Examples of distinct motion patterns of both IFT172 and KIF13B.** Cells expressing Halo-KIF13B (JF646) (red) and IFT172-eGFP (green) were imaged as described. Color overlay of time-lapse sequences of three different cells (A, C and E) are shown including, for the first two examples, the corresponding kymographs underneath (cell movement was too large to generate a kymograph reliably in the last case). Every third frame is shown (i.e., there are 1.5 sec between shown images). The first two examples illustrate the very different motion patterns for both proteins. Panel E shows additionally that IFT172-eGFP located in the tip of the cilium does not move together with Halo-KIF13B, which translocates from the upper to the distal part of the cilium (white arrows). Accumulation of Halo-KIF13B at the base is indicated with a blue arrow.

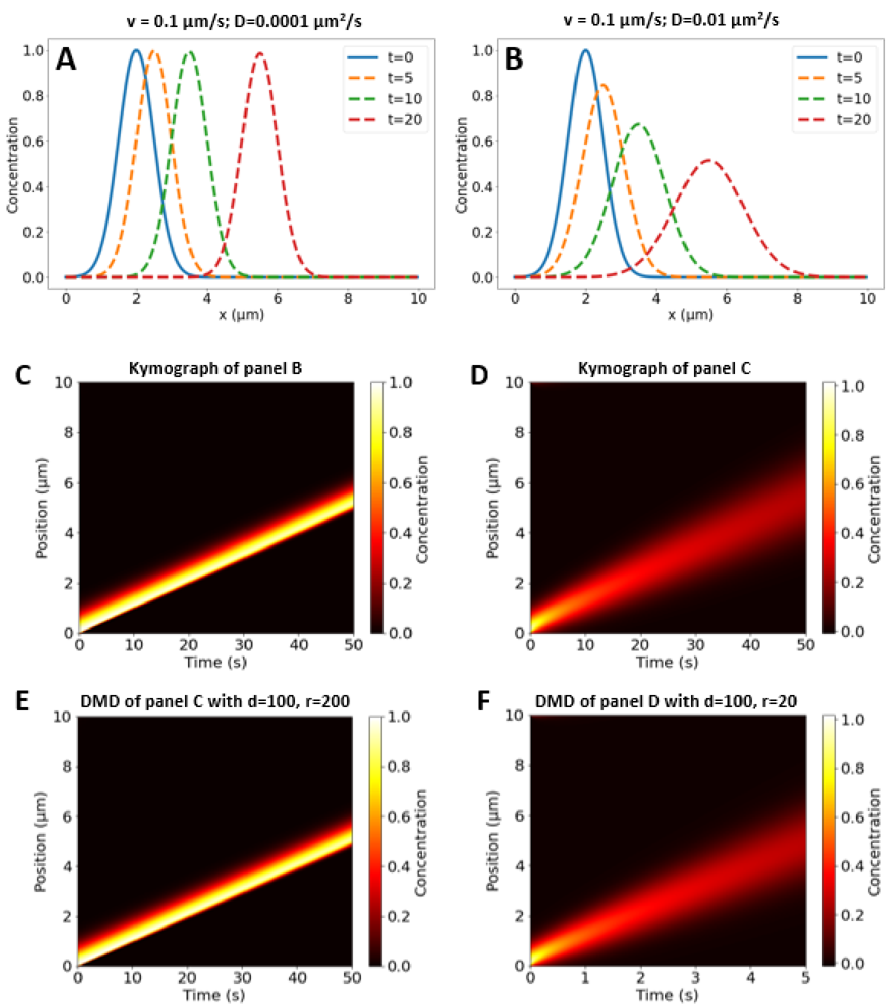


**Figure S6 Solution of the advection-diffusion equation and its reconstruction using DMD.** The solution of the advection-diffusion equation was plotted for the indicated parameters at selected time points for negligible (A) and significant diffusion (B). Advection causes translation to the right (i.e. into the cilium), while diffusion causes spreading of the protein. Both scenarios were also used to simulate kymograph (C and D), which were reconstructed by DMD with delay embedding using the indicated parameters (E and F).

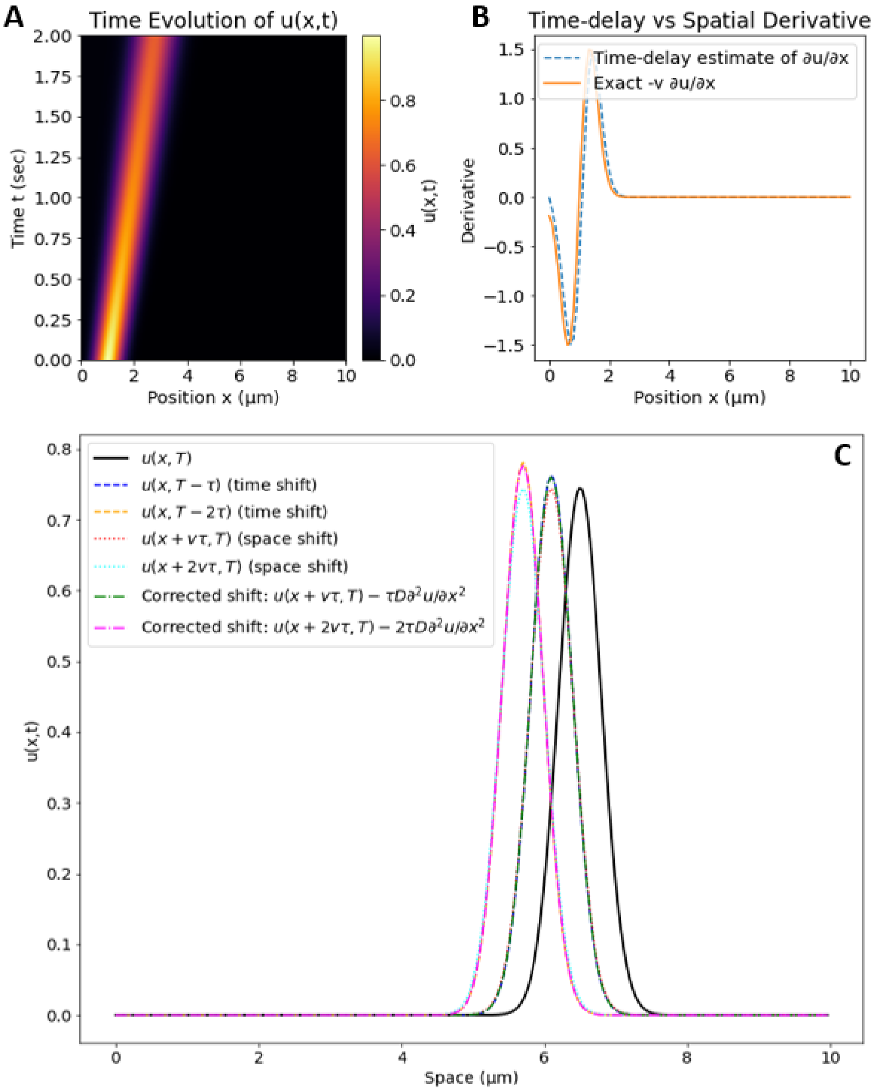


**Figure S7. Equivalence of space derivative and time-delay embedding as used in DMD.** Simulation of the advection-diffusion equation as in Fig. S5 (A) is shown next to the time-delay estimate of ∂u/∂x compared to the drift term of the advection-diffusion equation -v·∂u/∂x (B). The flow velocity was v = 2.0 µm/s and the diffusion constant, D = 0.01 $\mu m^2$/sec over a domain length of L = 10 µm, corresponding to a Peclet number of Pe = 0.0005, i.e. a process dominated by active transport. The negative of the spatial derivative multiplied with the velocity (orange line) was compared to the time-delay estimate (dashed blue line, B). Both methods were used to compare solutions of the advection-diffusion problem at selected time points (C). Here, the time-delayed version of the advection-diffusion solution ('time shift') is compared to the pure drift term ('space shift') or the drift + diffusion term ('corrected shift'). See supplemental text and appendix for further details.

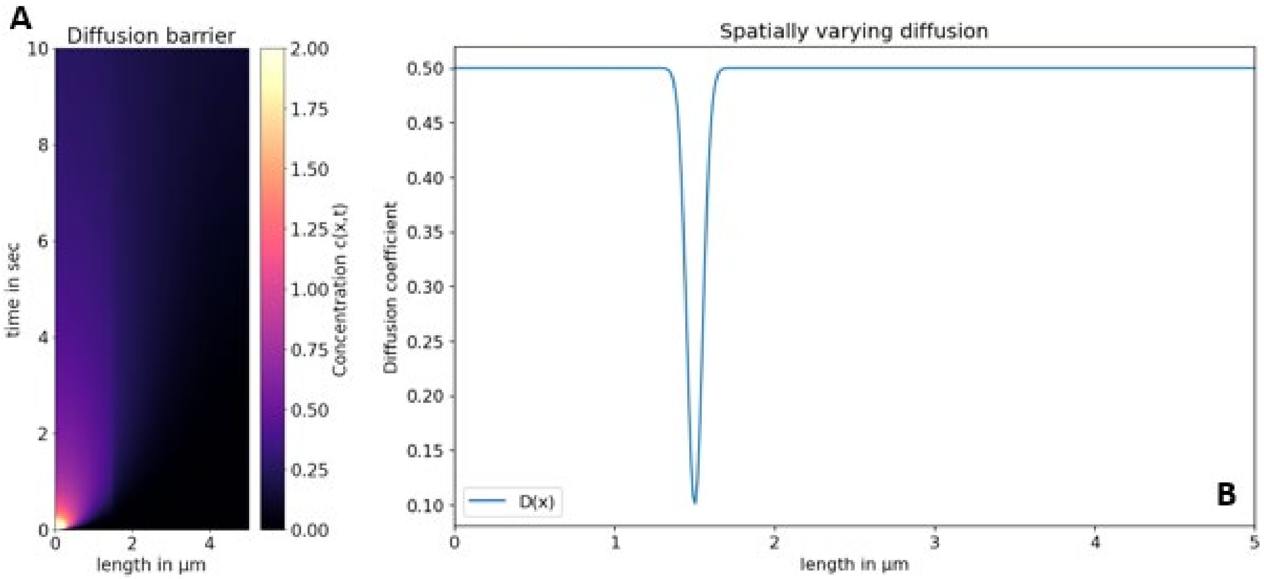


**Figure S8. Simulation of advection-diffusion inside cilia in the presence of a diffusion barrier.** Simulation of the advection-diffusion equation was carried out, as described above, but with a spatially varying diffusion constant, dropping from D = 0.5 µm$^2$/sec to 0.1 µm$^2$/sec at 1.5 µm distance from the starting point at the ciliary base. The drift velocity was set to v = 0.0 µm/sec, mimicking pure diffusion from the base into the cilium. The simulated concentration profile (A) and the space-dependent diffusion constant, $D(x)$, are shown (B). See Materials and methods for simulation details.

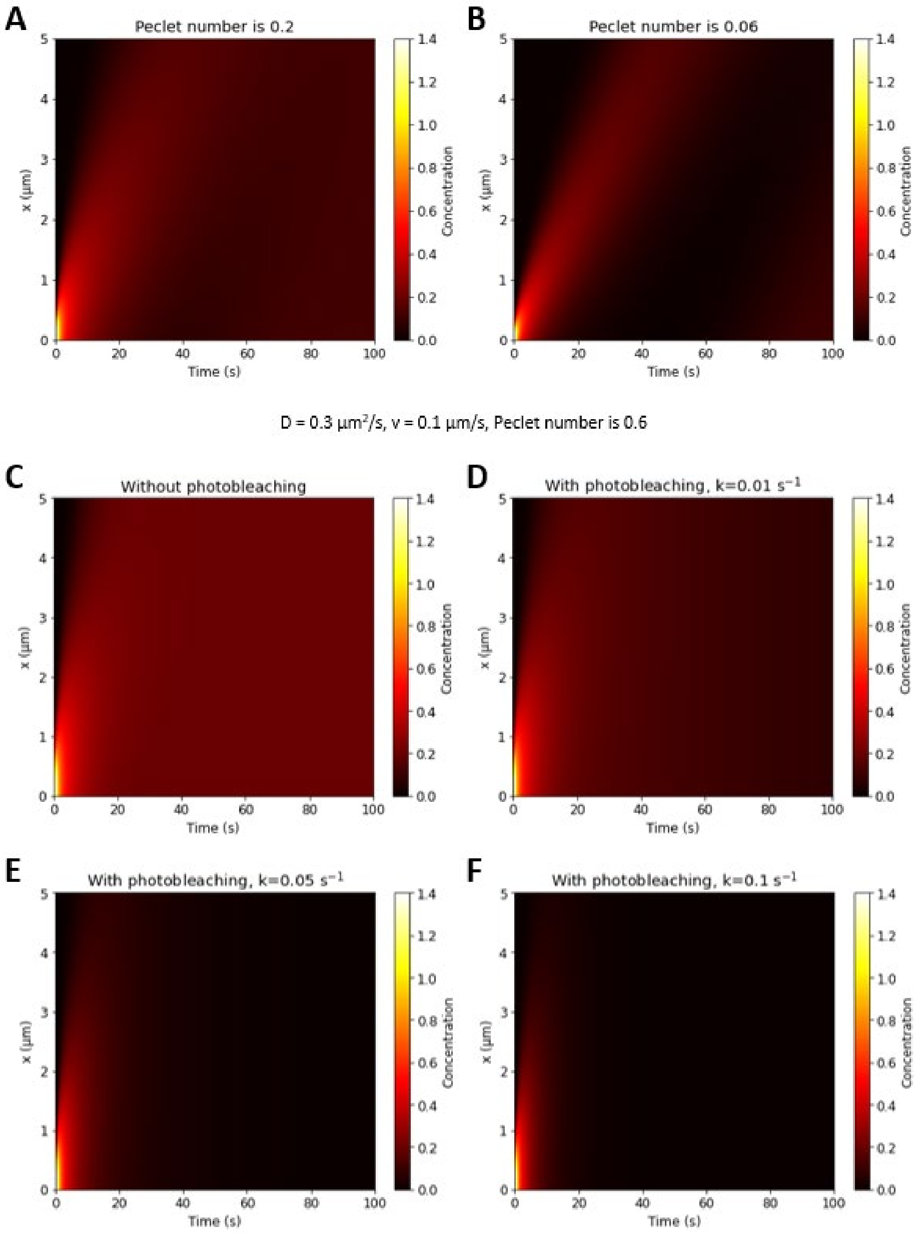


**Figure S9. Diffusion of fluorescent proteins can be masked by their photobleaching inside the cilium.** The impact of photo-bleaching on advection diffusion in kymo-graphs was simulated by plotting the solution of the corresponding advection-diffusion-reaction equation as function of space and time to emulate a kymograph, Eq. A16. The drift velocity was kept constant, while the diffusion constant was varied. In the absence of photobleaching higher Peclet numbers (Pe = 0.2, A) give a 'halo'-effect due to spreading by diffusion for long times compared to lower Peclet number (Pe = 0.06, B), where the trace of the simulated protein resembles a slightly spreading line. The effect of diffusion becomes even more pronounced for Pe = 0.6 in the absence of photobleaching (C), while slow but continuous photobleaching of the entire pool reduces the visible 'halo'-effect for long times (D). Under those conditions, the kymograph with high diffusion and photobleaching resembles that with lower diffusion and without photobleaching (compare D and A). Even stronger photobleaching can completely mask the diffusing component and also shorten the traces ( E and F).

**Table S1: reagents and resources used in this study**

| Reagent type (species) or resource | Designation | Source or reference | Identifiers | Additional information |
|---|---|---|---|---|
| Cell line (*Homo sapiens*) | hTERT-RPE1 expressing IFT172-eGFP | [1] | IFT172-eGFP | WT parental |
| Cell line (*H. sapiens)* | hTERT-RPE1 co-expressing IFT172-eGFP and Halo-KIF13B | This study | Pool | Generated by lentiviral transduction |
| Cell line (*H. sapiens*) | hTERT-RPE1 *KIF13B*$^{-/-}$ expressing IFT172-eGFP; Clone B3 | This study | Clone B3 | Generated by CRISPR/Cas9 methodology |
| Cell line (*H. sapiens*) | hTERT-RPE1 *KIF13B*$^{-/-}$ expressing IFT172-eGFP; Clone B4 | This study | Clone B4 | Generated by CRISPR/Cas9 methodology |
| Cell line (*H. sapiens*) | HEK293T | ATCC | Cat# CRL-3216 | |
| Strain, strain background (*Escherichia coli*) | DH10α | Lab stock | N/A | |
| Antibody | Anti-acetylated alpha-tubulin (mouse monoclonal) | Sigma-Aldrich | Cat# T7451 | IFM (1:2000) |
| Antibody | Anti-ARL13B (rabbit polyclonal) | Proteintech | Cat# 17711-1-AP | IFM (1:600) |
| Antibody | Anti-FBF1 (rabbit, polyclonal) | Proteintech | Cat# 11531-1-AP | IFM (1:600) |
| Antibody | Anti-GAPDH (rabbit polyclonal) | Cell Signalling Technology | Cat# 2118 | WB (1:1000) |
| Antibody | Anti-KIF13B (mouse monoclonal) | Merck | Cat# SAB1412812 | WB (1:600) |
| Antibody | Anti-Halo (rabbit polyclonal) | Promega | Cat# G9281 | WB (1:1000) |
| Antibody | Alexa Fluor™ 488 Donkey anti-Rabbit IgG (H+L) Highly Cross-Adsorbed Secondary Antibody | Invitrogen | Cat# A-21206 | IFM (1:600) |
| Antibody | Alexa Fluor™ 568 Donkey anti-Mouse IgG (H+L) Highly Cross-Adsorbed Secondary Antibody | Invitrogen | Cat# A10037 | IFM (1:600) |
| Antibody | Abberior STAR ORANGE, nanobody | Abberior | Cat# N2402-AbORANGE-S | IFM (1:500) |

| | | | | |
|---|---|---|---|---|
| | FluoTag®-X2, anto-Rabbit IgG | | | |
| Antibody | Polyclonal Goat Anti-Mouse Immunoglobulins/HRP | Dako | Cat# P0447 | WB (1:10000) |
| Antibody | Polyclonal Swine Anti-Rabbit Immunoglobulins/HRP | Dako | Cat# P0399 | WB (1:10000) |
| Chemical Reagent | DAPI | Sigma-Aldrich | Cat# D9542 | |
| Chemical Reagent | Immu-Mount | Thermo Fisher Scientific | Cat# 9990402 | |
| Chemical Reagent | Lipofectamine 3000 | Thermo Fisher Scientific | Cat# L3000015 | |
| Chemical reagent | SPY555-tubulin | Cytoskeleton, Inc. | Cat# CY-SC203 | Dissolved in DMSO and used at 1:1000 final concentration |
| Chemical Reagent | SiR-tubulin | Cytoskeleton, Inc. | Cat# CY-SC002 | Dissolved in DMSO and used at 1 mM final concentration |
| Chemical Reagent | Blasticidin S HCL | Gibco | Cat# r210-01 | |
| Chemical Reagent | Ciliobrevin D | Merck | Cat# 250401 | |
| Chemical Reagent | Forskolin | Merck | Cat# F6886 | |
| Chemical Reagent | Janelia Fluor® 646 HaloTag® Ligand | Promega | Cat# GA1120 | |
| Chemical Reagent | NuPAGE™ LDS Sample Buffer (4X) | Thermo Fischer Scientific | Cat# NP0007 | |
| Chemical Reagent | Penicillin-streptomycin | Sigma-Aldrich | Cat# P0781 | |
| Chemical Reagent | Trypsin MS approved | Serva | Cat# 37286.04 | |
| Chemical Reagent | M-PER Mammalian Protein Extraction Reagent | ThermoFisher Scientific | Cat# 78501 | |
| Commercial assay | Immobilon® -FL PVDF membrane | Sigma-Aldrich | Cat# 05317 | |
| Commercial assay | Mini-PROTEAN® TGX™ Precast Gel 4-15% 10 wells | Bio-Rab Laboratories | Cat# 456-1083 | |
| Commercial assay | Mini-PROTEAN® TGX™ Precast Gel 4-15% 15 wells | Bio-Rab Laboratories | Cat# 456-1086 | |

| | | | | |
|---|---|---|---|---|
| Commercial assay | PageRuler$^{TM}$ Plus Prestained Protein Ladder | Thermo Fischer Scientific | Cat#26619 | |
| Commercial assay | SuperSignal$^{TM}$ West Pico PLUS chemiluminescent Substrate | Thermo Fischer Scientific | Cat# 34580 | |
| Commercial assay | Trans-Blot® Turbo$^{TM}$ Transfer Pack, Mini format 0.2 µm Nitrocellulose membranes | Bio-Rad Laboratorie s | Cat# 1704158 | |
| Commercial assay | SpCas9 2NLS Nuclease | Synthego | | |
| Commercial assay | P3 Primary Cell 4D-Nucleofector® X Kit S | Lonza Bioscience | Cat# V4XP-3032 | |
| Commercial assay | NucleoBond Xtra Midi EF Kit | Macherey-Nagel | Cat# 740420.50 | |
| Sequence-based reagent | *H. sapiens Kif13b* exon 18 sgRNA | Synthego | sgRNA 1 | 5'-CGUGCUCAUACAUAAGCCUC -3' |
| Sequence-based reagent | *H. sapiens Kif13b* exon 18 sgRNA | Synthego | sgRNA 2 | 5'-UCAGCAACGCUUAAGACAGU -3' |
| Sequence-based reagent | *H. sapiens KIF13B* PCR oligo exon 17 | Eurofins genomics | | 5'-GTGTGTTAGAGGTGGCAAGC -3' |
| Sequence-based reagent | *H. sapiens KIF13B* PCR oligo exon 17 | Eurofins genomics | | 5'-CTGCAGGGAAGATGGTTGAG -3' |
| DNA plasmid | pENTR20-mNG-KIF13B-FL (Gateway) | | | Human KIF13B full-length was obtained by cloning from this plasmid. |
| DNA plasmid | pENTR20-Halo-C1 (Gateway) | Dr. Kay Oliver Schink, Oslo University Hospital, Norway | | |
| DNA plasmid | pCDH-EF1A-GW-IRES-blast (Gateway) | [2] | | |
| DNA plasmid | pMD2.G | Carlo Rivolta, Institute of Molecular and Clinical Ophthalmol ogy Basel, Switzerland | | |

| | | | | |
|---|---|---|---|---|
| DNA plasmid | pCMVΔ-R8.2 | Carlo Rivolta, Institute of Molecular and Clinical Ophthalmology Basel, Switzerland | | |
| DNA plasmid | Halo-KIF13B/ pCDH-EF1A-GW-IRES-blast | This study | LBP lab plasmid #1157 | Lentiviral plasmid coding for Halo-KIF13B |
| Other | 4D Nucleofector System X Unit | Lonza Bioscience | Cat# AAF-1003X | |
| Other | 4D Nucleofector System Core Unit | Lonza Bioscience | Cat# AAF-1003B | |
| Other | Trans-BLOT® Turbo Transfer System | Bio-Rad Laboratories | Cat# 1704150 | |
| Other | BioDrop µLITE | Biochrom | Cat# 4AJ-6319282 | |
| Software | Matlab R2024b | The MathWorks Inc | Version 24.2.0.2863752 (R2024b) Update 5 | |
| Software | Adobe Illustrator | Adobe | Version 29.1 | |
| Software | Adobe Photoshop | Adobe | Version 26.4.1 | |
| Software | Wolfram Mathematica | Wolfram Alpha LLC | Version 14.2 | |
| Software | KymoButler | Jakobs *et al.* [3] | | |
| Software | FIJI (Image J) | Schindelin *et al.* [4] | https://imagej.net/ij/ | |
| Software | GraphPad Prism | GraphPad Software Inc. | Version 10.4.2 | |

### Supporting video legends

**Movie 1. Intraciliary movement of IFT172-eGFP in parental hTERT-RPE1 cells starved for 24 hours. Related to Figure 1.** Representative video of non-treated parental cells stably expressing IFT172-eGFP. Cells were serum starved for 24 hours and stained for 1 hour with Spy555-tubulin (orange) to view the ciliary axoneme and microtubule cytoskeleton. Frame rate: 0.5 sec.

**Movie 2. Burst-like intraciliary movement of Halo-KIF13B in parental cells starved for 6 hours.** Representative live cell imaging of cells stably expressing IFT172-eGFP (green) and Halo-KIF13B (red). Cells were serum-starved for 6 hours. Transient accumulation of Halo-KIF13B was observed in 20 out of 67 cilia (29.8 %) imaged by this approach. Frame rate: 0.5 sec.

**Movie 3. Intraciliary movement of Halo-KIF13B in parental cells starved for 24 hours, with rapid expulsion of a vesicle-like particle from the tip. Related to Figure 3.** Live cell imaging of parental cells stably expressing IFT172-eGFP (green) and Halo-KIF13B (red). Cells were serum-starved for 24 hours. Expulsion of Halo-KIF13B-positive vesicle-like particles was observed in 2 out of 95 cilia (2.1 %) imaged by this approach. Frame rate: 0.5 sec.
